# Activating P2Y_1_ receptors improves function in arteries with repressed autophagy

**DOI:** 10.1101/2021.05.27.445652

**Authors:** Jae Min Cho, Seul-Ki Park, Oh Sung Kwon, D. Taylor La Salle, James Cerbie, Caitlin C. Fermoyle, David Morgan, Ashley Nelson, Amber Bledsoe, Leena P. Bharath, Megan Tandar, Satya P. Kunapuli, Russell S. Richardson, Pon Velayutham Anandh Babu, Sohom Mookherjee, Bellamkonda K. Kishore, Fei Wang, Tianxin Yang, Sihem Boudina, Joel D. Trinity, J. David Symons

## Abstract

**Objective:** The importance of endothelial cell (EC) autophagy to vascular homeostasis is evolving. Earlier we reported that purinergic 2Y_1_ receptor (P2Y_1_-R) activation rejuvenates shear-stress induced nitric oxide (NO) generation in bovine aortic endothelial cells that is otherwise compromised after pharmacological and genetic autophagy repression. Here we determined the translational and functional relevance of these findings.

**Approach and Results:** First we assessed translational relevance using older humans and mice that exhibit blunted EC autophagy at rest together with impaired arterial function vs. appropriate controls. Rhythmic handgrip exercise elevated radial artery shear rate similarly in adult and older males for 60-min. Compared to baseline, autophagy initiation, p-eNOS^S1177^ activation, and NO generation, occurred in radial artery ECs from adult but not older subjects. Regarding mice, indexes of autophagy and p-eNOS^S1177^ activation were robust in ECs from adult but not older mice in response to 60-min treadmill-running. Next we questioned whether an inability to initiate EC autophagy precipitates arterial dysfunction. Age-associated reductions in intraluminal flow-mediated vasodilation observed in older vs. adult mice were recapitulated in arteries from adult mice by : (i) NO synthase inhibition; (ii) autophagy impairment using 3-methyladenine (3-MA); (iii) EC Atg3 depletion (iecAtg3KO mice); (iv) P2Y_1_-R blockade; and (v) germline depletion of P2Y_1_-Rs. Importantly, P2Y_1_-R activation using 2-methylthio-ADP (2-Me-ADP) improved vasodilatory capacity in arteries from : (i) adult mice treated with 3-MA; (ii) adult iecAtg3KO mice; and (iii) older animals with repressed EC autophagy.

**Conclusions:** Arterial dysfunction concurrent with pharmacological, genetic, and age-associated EC autophagy compromise is improved by activating P2Y_1_-Rs.

## Introduction

Autophagy is critical for maintaining quality control of the cell^1^ and integrity of the cardiovascular system.^2^ Aging compromises the process of autophagy in a number of cell types.^3, 4^ With specific regard to endothelial cells (ECs), one report indicates a protein responsible for autophagy initiation was lower (i.e., beclin-1), and a marker of undegraded autophagy substrates was higher (i.e., p62/sequestosome-1), in primary brachial artery ECs from older vs. adult subjects under basal conditions.^5^ Because older participants in that study displayed brachial artery NO-dependent endothelial dysfunction, the authors suggested a link might exist between defective EC autophagy and compromised EC NO generation.

Since the circulating milieu and EC phenotype associated with aging is complex, we used a reductionist approach to determine whether EC autophagy suppression *per se* is sufficient to impair shear-stress induced NO generation in bovine aortic ECs (BAECs). Inhibiting EC autophagy via pharmacological and genetic approaches prevented shear stress-induced activating phosphorylation of eNOS at serine 1177 (p-eNOS^S1177^) and NO generation, and heightened shear stress-evoked reactive oxygen species (ROS) production and pro-inflammatory gene expression.^6^ We concluded that EC autophagy plays a critical role in maintaining NO bioavailability, and contributes importantly to oxidant / antioxidant and inflammatory / anti-inflammatory balance in ECs. After recapitulating these findings in human arterial endothelial cells (HAECs), we revealed that genetic repression of autophagy impairs EC glycolysis and subsequent autocrine signaling via the purinergic 2Y_1_ receptor (P2Y_1_-R) to eNOS, to an extent that NO generation is compromised ^7^. By manipulating P2Y_1_-R signaling this phenotype was repeated in ECs with intact autophagy, and rescued in ECs with genetic repression of autophagy.^7^ Here we sought to determine the translational and functional relevance of these findings by addressing three questions. First, is autophagy and NO generation impaired in arterial ECs from older vs. adult humans and mice in response to shear-stress evoked by functional hyperemia? Second, does repressed shear-stress induced NO generation after autophagy diminution *in vitro* translate to impaired intraluminal flow-mediated arterial vasodilation examined *ex vivo*? If so (i.e., third), can limited intraluminal flow-mediated arterial vasodilation in the context of pharmacological, genetic, and aging-associated EC autophagy compromise in mice be rejuvenated by targeting P2Y_1_-Rs?

Here we report that elevated arterial shear-rate associated with active hyperemia evoked by rhythmic handgrip exercise initiates autophagy, p-eNOS^S1177^ activation, and NO generation in primary arterial ECs obtained via j-wire from the radial artery of adult but not older participants. Further, robust activation of arterial EC autophagy and NO generation in response to an acute bout of treadmill-running was observed in adult but not older mice. These results suggest our earlier findings that shear-stress-induced NO generation is prevented by pharmacological and genetic autophagy repression in BAECs and HAECs can be translated to primary arterial ECs from older vs. adult humans and mice. Next we demonstrated that P2Y_1_-R activation rejuvenates intraluminal flow mediated vasodilation that is otherwise attenuated in femoral arteries from : (i) adult mice (7-month) after pharmacological autophagy compromise; (ii) adult mice (4-month) with inducible depletion of Atg3 specifically in ECs; and (iii) older mice (24-month) that display indexes of repressed arterial EC autophagy at baseline and in response to 60-min treadmill-running. Collectively, solid evidence is provided that targeting purinergic signaling to endothelial NO synthase in the context of repressed vascular autophagy mitigates arterial dysfunction.

## Materials and Methods

Protocol approval and written informed consent for studies involving humans was obtained according to the University of Utah (UU) and the Department of Veterans Affairs Salt Lake City Health Care System Institutional Review Boards. Adult and older male subjects visited the laboratory on two occasions separated by at least 7 days. On day 1 a flow-mediated dilation (FMD) test was completed and a maximal handgrip workload evaluation was performed ^8^. On day 2 radial artery ECs were collected via J-wire before and after rhythmic handgrip exercise (RHE) that elevated arterial shear-rate for 60-min.^8^ Protocols approved by UU Institutional Animal Care and Use Committee were completed on male : (i) adult and older mice; (ii) mice with tamoxifen-inducible depletion of Atg3 specifically in ECs and their wild-type littermates; and (iii) mice with germline depletion of P2Y_1_-Rs and their wild-type littermates. Detailed procedures using humans, mice, and cell-systems assays, together with a comprehensive listing of all reagents in the *Major Resource Table,* are located in the Supplemental Methods section of the Data Supplement.

### Statistics

Information concerning statistical tests is provided for each data set in the respective figure or table legend.

## Results

### ECs from older males with impaired arterial function are resistant to elevated arterial shear-rate concerning autophagy initiation and NO generation

We reported that elevated arterial shear-rate associated with functional hyperemia evoked by rhythmic handgrip exercise (RHE) increases autophagy initiation, and NO and O_2_^•-^ production, in radial artery ECs obtained from healthy adult males ^8^. These findings, coupled with our earlier results that pharmacologic and genetic repression of EC autophagy prevents shear-stress induced NO generation in HAECs and BAECs, prompted us to test the hypothesis that shear-induced autophagy initiation and NO generation are blunted in ECs from older vs. adult volunteers **Table I in the Data Supplement**).

As anticipated, flow-mediated dilation peak was lower in older vs. adult subjects (**Figure 1A**). Comparing RHE-Pre between groups i.e., the influence of aging, p-eNOS^S1177^ and NO generation were lower (**Figure 1B through 1D**) and O_2_^•-^ was higher (**Figure 1B and 1E**) in ECs from older vs. adult participants, whereas no differences concerning total eNOS protein expression existed between groups (**Figure I in the Data Supplement**). Regarding autophagy, while microtubule-associated proteins 1A/1B light chain 3B (LC3B), LC3B-bound puncta, and LC3B colocalization with lysosomal associated membrane protein 2a (LAMP2A) were lower in ECs from older vs. adult participants under baseline conditions (i.e., RHE-Pre), we observed no differences between adult and older subjects concerning autophagy related gene 3 (Atg3), Beclin-1, or p62 (**Figure 2**).

**Figure 1.**
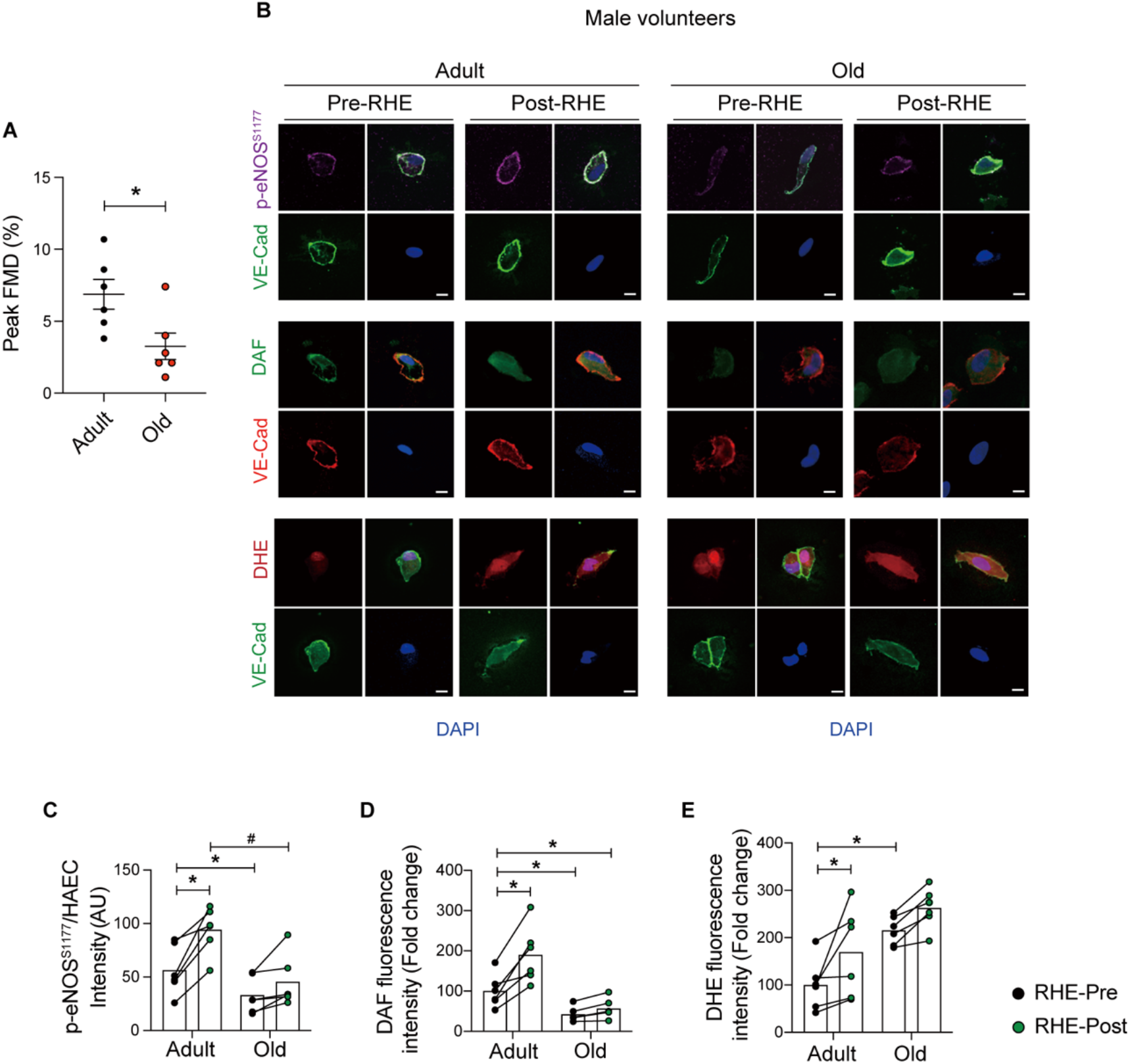
Endothelial cells from older males with impaired arterial function are resistant to elevated radial artery shear-rate concerning nitric oxide generation and reactive oxygen species. (**A**) Flow-mediated dilation (FMD, %) peak was lower in older vs. adult male subjects. Representative images (**B**) and mean immunofluorescence staining intensity (**C**-**E**) is shown. Comparing RHE-Pre to RHE-Pre between groups i.e., the influence of aging, p-eNOS^S1177^ and DAF are lower in ECs from older vs. adult participants, while DHE staining is elevated in ECs from older vs. adult subjects (i.e., histogram 1 vs. 3). Comparing RHE-Pre to RHE-Post between groups i.e., the influence of elevated arterial shear rate, p-eNOS^S1177^, DAF, and DHE are elevated in ECs from adult (bar 1 vs. 2) but not older (bar 3 vs. 4) subjects. Seventy-five ECs from each time point and each subject were measured, and the staining intensity was normalized to values obtained from commercial HAECs using identical conditions. Data are from the same 6 subjects per group as shown in *A*. *p<0.05 vs RHE-Pre adult; #p<0.05 vs RHE-Post adult. Scale bar : 10 μm. Magnification : 60 X. Significance was assessed using an unpaired t-test (**A**) and a two-way ANOVA (**C**-**E**).

**Figure 2.**
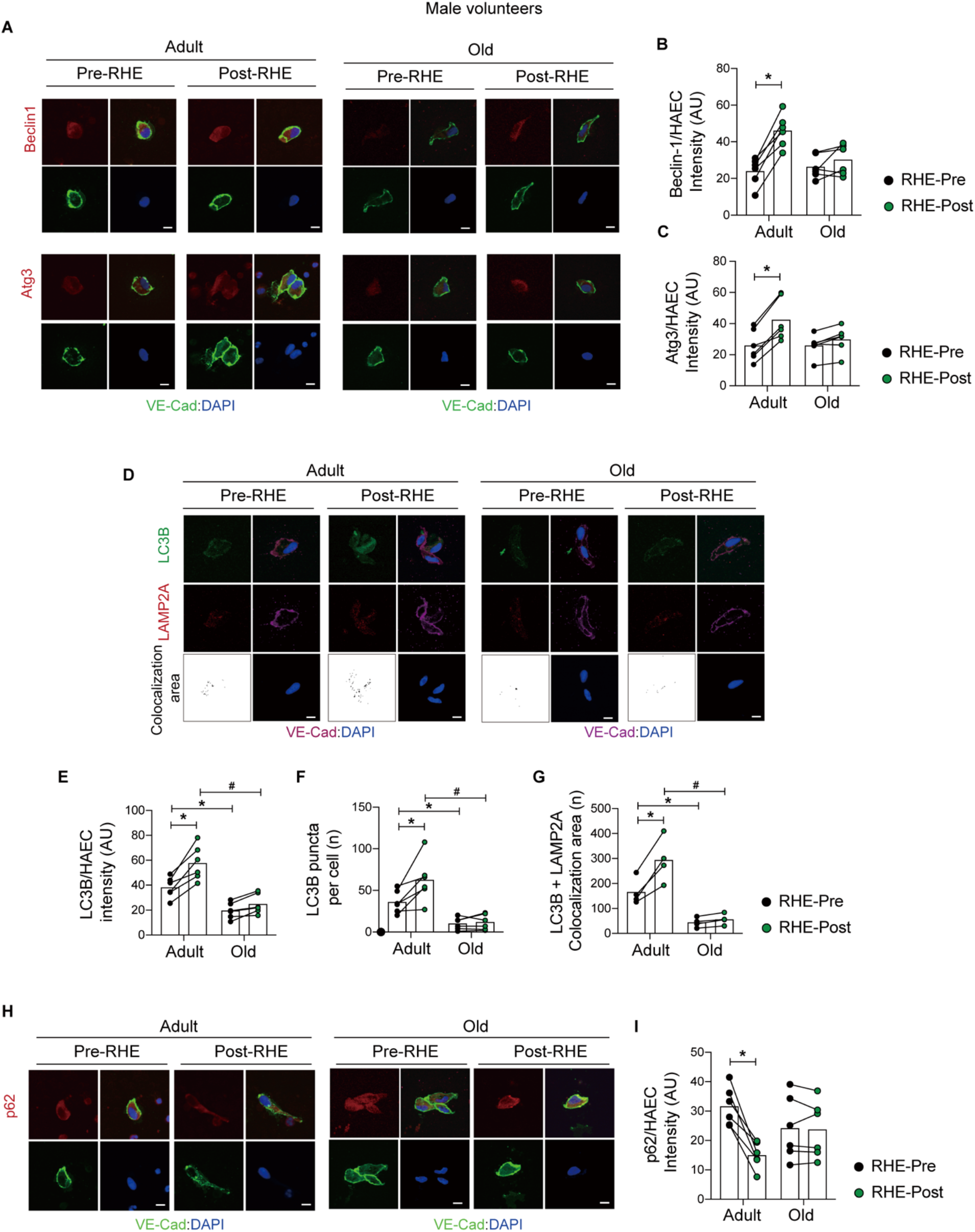
Endothelial cells from older males are resistant to elevated radial artery shear-rate concerning autophagy. Representative images (**A**, **D**, and **H**) and mean immunofluorescence staining intensity (**B**, **C**, **E-G**, and **I**) is shown. Comparing RHE-Pre to RHE-Pre between groups i.e., the influence of aging, LC3B expression, LC3B bound puncta, and LC3B colocalization with LAMP2A are lower in ECs from older vs. adult subjects, whereas Beclin-1, Atg3, and p62 are similar between groups (histogram 1 vs. 3). Comparing RHE-Pre to RHE-Post between groups i.e., the influence of elevated arterial shear rate, Beclin-1, Atg3, LC3B, LC3B bound puncta, LC3B colocalization with LAMP2A, and p62 degradation increased in ECs from adult (histogram 1 vs. 2) but not older (histogram 3 vs. 4) subjects. Seventy-five ECs from each time point and each subject were measured, and the staining intensity was normalized to values obtained from commercial HAECs using identical conditions. *p<0.05 vs RHE-Pre adult; #p<0.05 vs RHE-Post adult. Scale bar : 10 μm. Magnification : 60 X. Statistical significance was assessed using a two-way ANOVA (**B**, **C**, **E-G**, and **I**).

Relative to measures obtained at baseline i.e., RHE-Pre, exercise elevated brachial artery (BA) blood flow velocity and arterial shear rate ~ 3-fold for 60-min to values that were not different between adult and older subjects (**Movie I and Online Figure II in the Data Supplement**), and systemic hemodynamics were unaltered by this stimulus in both groups (**Table II in the Data Supplement**). When compared to values obtained at RHE-Pre, RHE-Post ECs displayed increased p-eNOS^S1177^, NO generation, and O_2_^•-^ production, in adult but not older subjects (**Figure 1 and Figure I in the Data Supplement**). Regarding autophagy, compared to RHE-Pre, increased expression of Beclin-1, Atg3, LC3B, LC3B+LAMP2A colocalization, LC3B bound puncta, and decreased expression (i.e., increased degradation) of p62, was displayed by ECs from adult but not older participants at RHE-Post (**Figure 2**). Providing translational relevance of our *in vitro* findings that pharmacologic and genetic repression of EC autophagy limits shear-stress induced EC NO generation, autophagy initiation and NO generation are blunted in ECs from older vs. adult males in response to elevated arterial shear rate associated with active hyperemia.

### Repressed EC autophagy in older vs. adult mice is associated with compromised intraluminal flow-mediated vasodilation

We sought to determine whether EC autophagy and intraluminal flow-mediated vasodilation are repressed in similar sized arteries in an aging-related manner. mRNA expression was assessed in ECs and media + adventitia (M+A) obtained from iliac arteries. Purity of the EC and M+A fraction was verified by quantifying mRNA expression of platelet/endothelial cell adhesion molecule 1 (*Pecam1*) and alpha-smooth muscle actin (*α-Sma*), respectively. *Pecam1* mRNA was highly expressed in ECs, but not in M+A, whereas *α-Sma* mRNA was highly expressed in M+A, but not in the EC fraction (**Figure 3A**). Using the EC fraction from iliac arteries, we demonstrate that mRNA expression of transcriptional factor EB (*Tfeb*), *Atg3, Atg5*, and *Map1lc3b* is lower in older vs. adult mice (**Figure 3B**). These findings in ECs from iliac arteries were recapitulated in ECs from carotid arteries from the same mice (**Figure III in the Data Supplement**).

**Figure 3.**
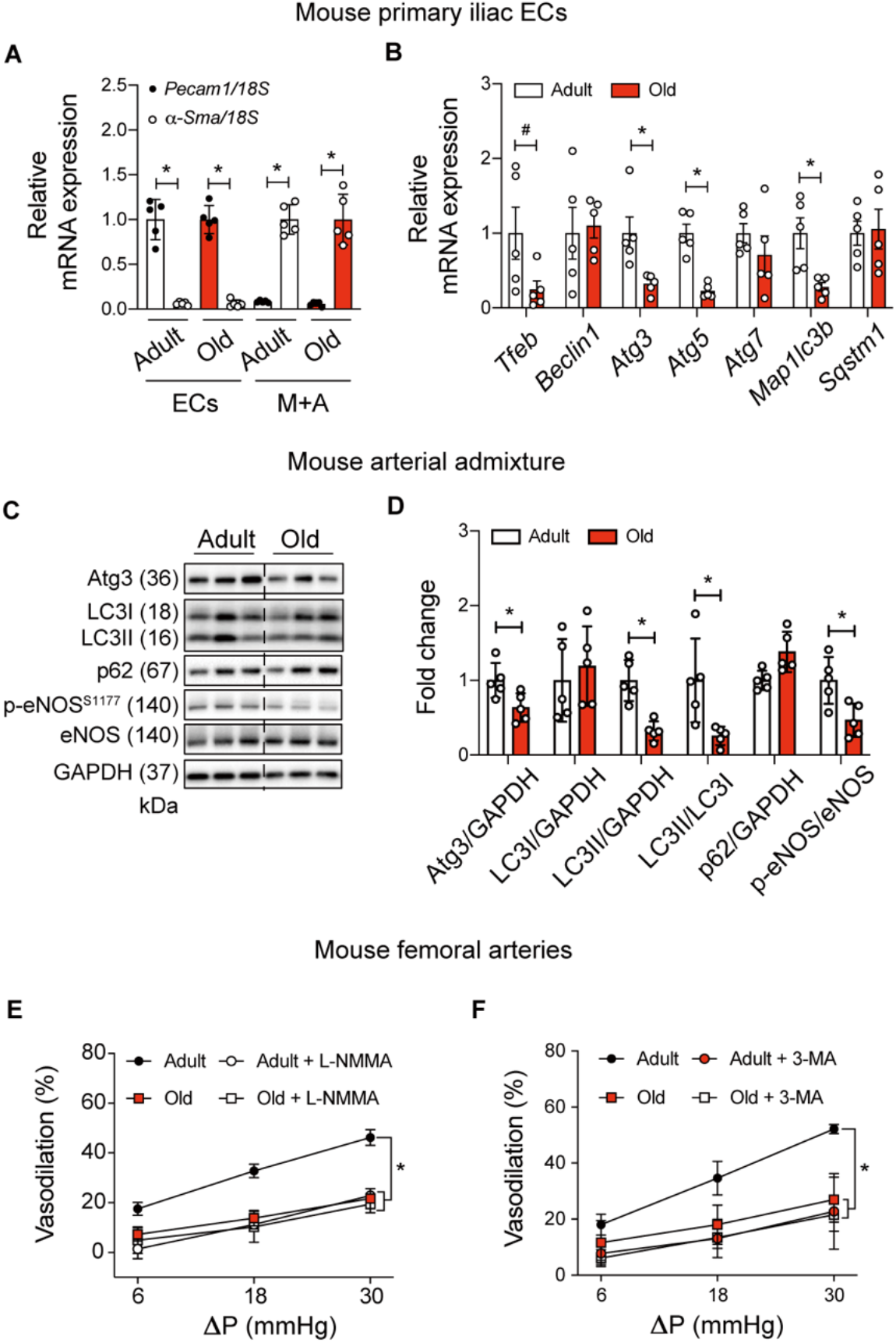
Repressed endothelial cell autophagy in older vs. adult mice is associated with compromised intraluminal flow-mediated vasodilation. *Pecam1* mRNA expression is robust whereas *α-Sma* is minimal in the EC fraction from iliac (**A**) arteries of adult and old mice. *α-Sma* mRNA expression is robust whereas *Pecam1* is minimal in the media + adventitia (M+A) fraction from iliac (**A**) arteries of adult and old mice. Iliac artery ECs (**B**) of older mice display repressed mRNA expression of autophagy indices vs. adult mice. For **A** and **B** all values are normalized by *18S*; n=5 mice per group x 2 iliac arteries per mouse. For **A**, *p<0.05 vs *Pecam1.* For **B** and **D**, *p<0.05 vs Adult. Representative images (**C**) and mean densitometry (**D**) from arterial admixtures indicate indexes of autophagy (normalized to GAPDH) and p-eNOS^S1177^ (normalized to eNOS) are compromised in older vs. adult mice. For **C** and **D**, n=5 per group; p<0.05 vs Adult. (**E**) Intraluminal flow-mediated vasodilation (%) was : (i) greater in femoral arteries from adult vs. older mice; and (ii) sensitive to L-NMMA in femoral arteries from adult but not older mice. Robust intraluminal flow-mediated vasodilation is displayed by arteries from adult but not older mice in the presence of vehicle (DMSO, **F**). After a 30-min incubation with : (i) 5 mmol/L 3-MA, intraluminal flow-mediated vasodilation is repressed in arteries from adult but not older mice (**F**). For **E** and **F**, n=6 mice per group x 1 artery per mouse. *p<0.05 vs adult. Statistical significance was assessed by unpaired t-test (**A**-**D**) and a two-way repeated measures ANOVA (**E** and **F**).

To obtain sufficient substrate for protein determination, admixtures of aorta, iliac, and femoral artery were used. Protein expression of Atg3, the ratio of the lipidated form of LC3-II to GAPDH (LC3-II:GAPDH), LC3-II:LC3-I, and p-eNOS^S1177^:eNOS, were lower, whereas expression of p62 trended higher (p=0.09), in arteries from older vs adult mice (**Figure 3C and 3D**). These findings collectively support that indexes of autophagy-related mRNA and protein expression are lower in ECs and arterial lysates from old vs adult mice.

Next we tested whether impaired NO-mediated vasodilation exists in femoral arteries from old mice with repressed EC autophagy vs. adult mice with intact EC autophagy. Animal and vessel characteristics are shown in **Table III and Figure IV in the Data Supplement**. The design of this and subsequent vascular function experiments required that two intraluminal flow-mediated vasodilation curves be performed using the same artery. Because of this, we confirmed in preliminary experiments that : (i) pressure gradients (ΔP) of 6, 18, and 30 mmHg evoke similar flow-rates when separated by 30-min; (ii) ΔPs of 6, 18, and 30 mmHg cause vasodilatory responses that are not different when separated by 30-min; and (iii) phenylephrine (PE) -induced precontraction can be sustained (**Figure V in the Data Supplement**). Our findings, as anticipated based on the available literature, indicate that : (i) blunted vasodilation in response to three different flow rates exists in arteries from old vs. adult mice; (ii) intraluminal incubation with 10^-3^ mol/L N^G^-monomethyl-L-arginine (L-NMMA) represses flow-mediated vasodilation in arteries from adult but not older mice (**Figure 3E**); and (iii) vasodilation in response to sodium nitroprusside is not different in arteries from adult and old mice (**Figure VIA in the Data Supplement**). These findings are the first to reveal that aging impairs intraluminal flow-mediated vasodilation in a NO-dependent manner in arteries wherein repressed EC and vascular autophagy have been documented.

### Shear stress -induced eNOS activation in HAECs and intraluminal flow-mediated vasodilation in femoral arteries are attenuated by 3-MA

Thus far, our results indicate an association exists between suppressed EC autophagy (**Figure 3B**) and impaired NO-mediated intraluminal flow mediated vasodilation in the context of aging (**Figure 3E**). However, because the circulating environment accompanying aging is complex, we sought to determine the independent influence from repressed autophagy to arterial dysfunction. First we verified that shear-stress - induced Atg3:GAPDH and LC3II:GAPDH expression, and p62:GAPDH degradation (**Figure VIIA through VIID in the Data Supplement**) are prevented by 3-MA and that this intervention limits p-eNOS^S1177^ : eNOS and NO generation (SI Appendix, Figure S8 A-D) in the absence of cell death or apoptosis (**Figure VIIE through VIIG in the Data Supplement**). Next we tested the hypothesis that arteries from adult mice treated with 3-MA mimic an aging phenotype. Supporting this, intraluminal flow-mediated vasodilation displayed by arteries from adult mice was blunted after autophagy inhibition using 3-MA, to an extent that was not different from responses exhibited by arteries from old mice in the absence (i.e., first response) or presence (i.e., second response, 30-min later) of 3-MA (**Figure 3F**). These data illustrate repressed arterial autophagy *per se* contributes to impaired arterial vasodilatory capacity.

### Treadmill-running activates EC autophagy and eNOS in arteries from adult but not older mice

Data shown in Figures 1 and 2 indicate autophagy initiation and NO generation are blunted in ECs from older vs. adult males that complete 60-min rhythmic handgrip exercise. Results in Figure 3B indicate an age-associated repression of arterial EC autophagy exists under basal conditions in mice. Here we determined whether autophagy and eNOS activation are impaired in arterial ECs from older vs. adult mice in response to active hyperemia evoked by 60-min treadmill-running. The mode, intensity, and duration of exercise was chosen based on its demonstrated ability to heighten arterial p-eNOS^S1177^ and eNOS enzyme activity in mice.^9^ Substantiating our hypothesis, treadmill-running increased expression of Atg3, LC3B, and p-eNOS^S1177^ to a lesser extent in aortic ECs from older vs. adult mice (**Figure 4**). These results are congruent with those observed in ECs from older vs. adult male participants (**Figure 1 and 2**).

**Figure 4.**
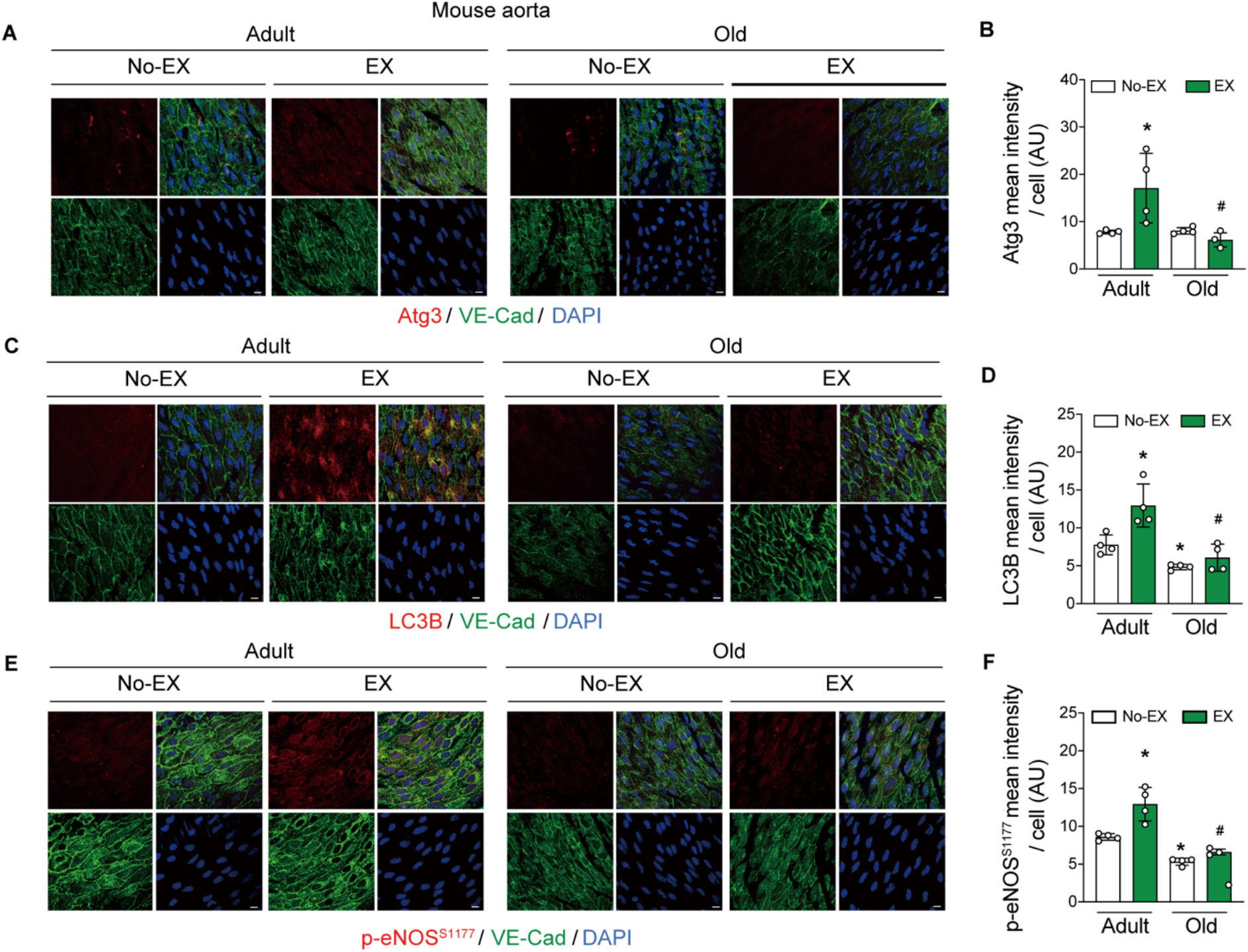
Endothelial cells from adult but not older mice display autophagy initiation and nitric oxide generation in response to 60-min treadmill running. Representative images (**A**, **C**, and **E**) and mean immunofluorescence data (**B**, **D**, and **F**) from aortic endothelial cells from mice that did not (No-EX) or did (EX) complete 60-min treadmill-running are shown. The four quadrants of each staining protocol represent VE-Cadherin (green, bottom left); DAPI (blue, bottom right); protein of interest (red, top left), and merge (yellow, top right). Atg3, LC3B, and p-eNOS^S1177^ (**B**, **D**, and **F**) increased in EX vs. No-EX adult (histogram 1 vs. 2) but not older (histogram 3 vs. 4) mice. Comparing No-EX to No-EX between groups i.e., the influence of aging (histogram 1 vs. 3), LC3B (**D**) and p-eNOS^S1177^ (**F**) were lower in older vs. adult mice, whereas Atg3 was not different between groups. Four mice per group, 5 images per aorta per mouse, and 20 ECs per image were quantified. * p<0.05 vs No-EX-Adult; # p<0.05 vs EX-Adult. Scale bar : 10 μm. Magnification : 60 X. Statistical significance was assessed using a one-way ANOVA.

### Attenuated shear-induced eNOS activation after 3-MA is restored by 2-Me-ADP in HAECs

Findings shown in **Figures VII and VIII in the Data Supplement** illustrate that 3-MA impairs shear-stress -induced autophagy and NO generation in HAECs. In agreement with our previous findings in BAECs ^7^, repressed shear-induced p-eNOS^S1177^ and NO generation after 3-MA was normalized in HAECs by concurrent treatment with the purinergic 2Y_1_ receptor (P2Y_1_-R) agonist 2-Me-ADP (**Figure VIII and IX in the Data Supplement**).

### Intraluminal flow-mediated vasodilation is impaired by aging and 3-MA, but is improved by 2-Me-ADP

Because 2-Me-ADP restored shear-stress induced p-eNOS^S1177^ and NO generation in HAECs after inhibiting autophagy initiation using 3-MA (**Figure VIII and IX in the Data Supplement**), we sought to determine whether a similar pattern of results is translated to arteries that display an aging-associated reduction in vascular autophagy. In this regard, intraluminal flow-mediated vasodilation was impaired in arteries from old vs. adult mice, but 2-Me-ADP rejuvenated vasodilatory responses in arteries from aged animals to values that were not different from adult mice treated with 2-Me-ADP (**Figure 5A**).

**Figure 5.**
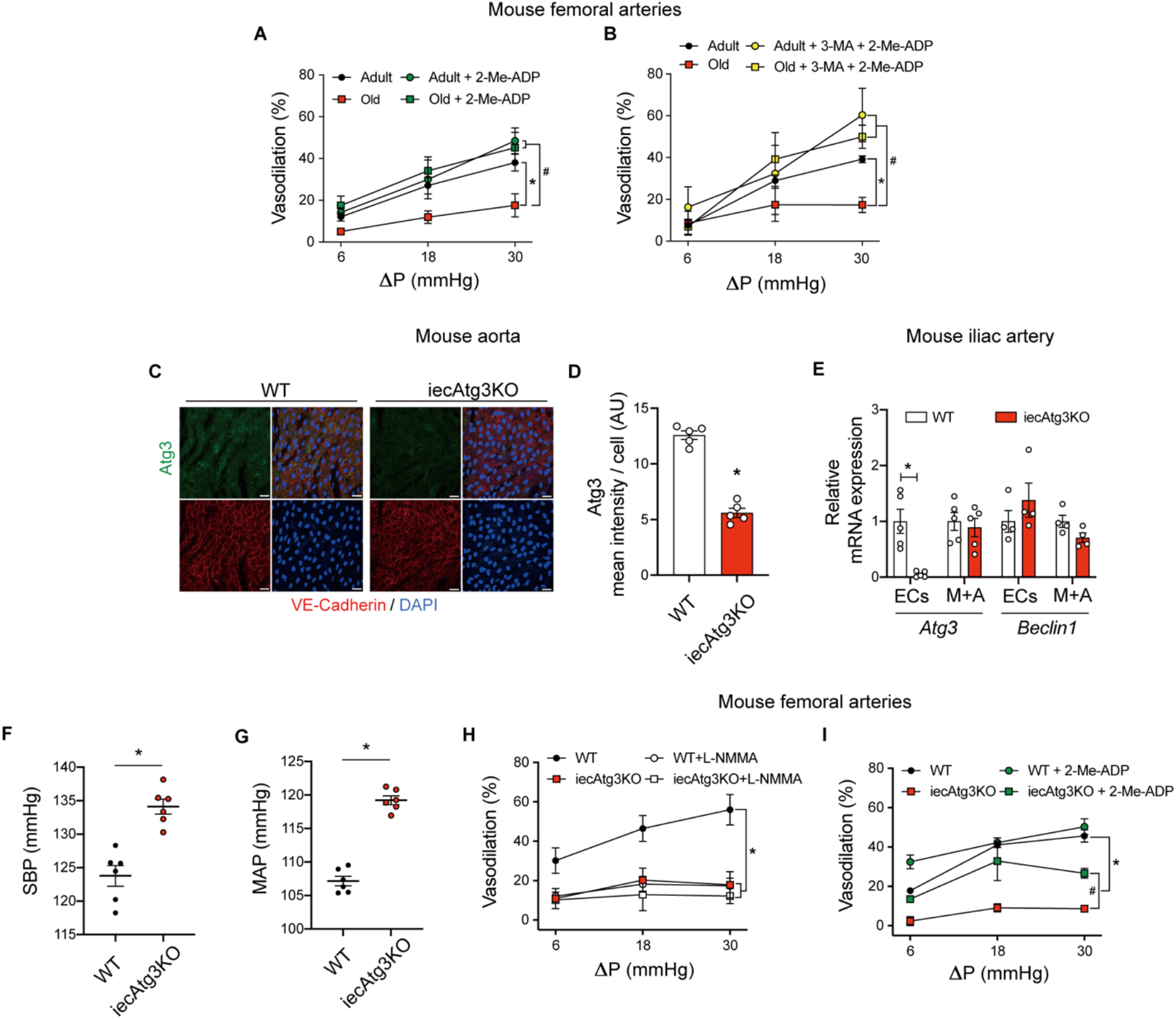
Impaired flow-mediated vasodilation evoked by pharmacological, 3-MA or genetic, iecAtg3KO in arteries from adult mice is reestablished by 2-Me-ADP. Robust intraluminal flow-mediated vasodilation is displayed by arteries from adult but not older mice in the presence of vehicle (DMSO; **A** and **B**). After a 30-min incubation with : (i) 100 μmol/L 2-Me-ADP, intraluminal flow-mediated vasodilation is improved in arteries from older mice (**A**); (ii) 3-MA + 2-Me-ADP, intraluminal flow-mediated vasodilation is improved in arteries from adult and older mice (**B**). For **A** and **B**, n=4 mice per group; One artery per mouse was used. For **A** and **B**, *p<0.05 vs Adult; #p<0.05 vs Old. Representative en face images (**C**) and mean immunofluorescent staining intensity (normalized by the number of cells; (**D**) of endothelial cells (ECs) from aorta of wild type (WT) mice and mice with inducible depletion of Atg3 in ECs. ECs were identified by co-staining for DAPI and VE-cadherin. For each quadrant (**C**): bottom right DAPI (blue); bottom left, VE-cadherin (red); top left: Atg3 (green); top right, merge. Atg3 (**B**) was lower in ECs from iecAtg3KO vs. WT mice. For *C* and *D*, n=3 segments of aorta from 4 mice in each group. For *C*, scale bar represents 10 μm, magnification=60X. (**E**) EC *Atg3* mRNA expression is minimal in iecAtg3KO vs. WT mice, whereas M+A *Atg3* mRNA expression is similar between groups. *Beclin1* mRNA expression is not different between groups in ECs or M+A. These data indicate knockdown of *Atg3* is robust, specific to ECs, and that other indexes of autophagy e.g., *Beclin1* are unaffected. Data shown in *E* were normalized by *18S;* n=5 mice per group x 2 iliac arteries per mouse. For **E**, *p<0.05 vs WT. *p<0.05 vs WT. Systolic blood pressure (SBP) and mean arterial pressure (MAP) were assessed via untethered biotelemetry. SBP (**F**) and MAP (**G**) data represent 3 x 24 h periods obtained 6-days post-surgery to implant the telemetry devices. For **F** and **G**, n=6 per group. *p<0.05 vs WT. Robust intraluminal flow-mediated vasodilation is displayed by femoral arteries from WT but not iecAtg3KO mice in the presence (**H** and **I**). After a 30-min incubation with : (i) 10^-3^ mol/L L-NMMA, intraluminal flow-mediated vasodilation is repressed in arteries from WT but not iecAtg3KO mice (**H**); (ii) 100 μmol/L 2-Me-ADP, intraluminal flow-mediated vasodilation is improved in arteries from iecAtg3KO mice (**I**). For **H** and **I**, n=5 mice per group, 1 artery per mouse. *p<0.05 vs WT; #p<0.05 vs iecAtg3KO. Statistical significance was assessed using a two-way repeated measures ANOVA (**A**, **B**, **H**, and **I**) and unpaired t-test (**C***-***G**).

Next we determined whether P2Y_1_-R activation reestablishes arterial function in a manner that is “downstream” from defective autophagy. Depressed intraluminal flow-mediated vasodilation was confirmed in arteries from older vs. adult mice. However, simultaneous incubation with 3-MA + 2-Me-ADP improved vasodilatory capacity in arteries from older mice to an extent that was not different from adult mice (**Figure 5B**). Importantly, neither 3-MA, 2-Me-ADP, nor their combination, influenced responses to sodium nitroprusside (**Figure VB through VD in the Data Supplement**), indicating vascular smooth muscle function was not affected by the respective treatments. These findings illustrate that purinergic reactivation of autocrine signaling using 2-Me-ADP improves intraluminal flow-induced vasodilation that is otherwise impaired in arteries with physiological (i.e., aging) or pharmacological (i.e., 3-MA) autophagy disruption. Animal and vessel characteristics are shown in **Table IV in the Data Supplement**. *Shear stress-induced eNOS activation and NO generation in HAECs is prevented by Atg3 knockdown but is restored by 2-Me-ADP.* Using a complementary approach to pharmacological autophagy repression, we knocked down Atg3 in HAECs using CRISPR-Cas9. Twenty dyn/cm^2^ shear-stress increased Atg3:GAPDH and LC3II:GAPDH expression, and p62:GAPDH degradation, in WT but not sgAtg3KO HAECs (**Figure XA through XD in the Data Supplement**). As expected, genetic diminution of Atg3 in HAECs prevented shear-induced eNOS activation and NO generation, but both endpoints were rejuvenated by concurrent treatment with 2-Me-ADP (**Figure XI in the Data Supplement**). None of the treatments influenced cell death or apoptosis (**Figure XI in the Data Supplement**).

### Mice with tamoxifen-inducible endothelial cell specific Atg3 depletion

To determine whether our findings from pharmacological and genetic autophagy repression in HAECs can be translated to intact animals, we generated mice with tamoxifen-inducible depletion of Atg3 specifically in ECs i.e., iecAtg3KO mice. Genotyping results and the tamoxifen regimen to induce depletion are shown in **Figure XIIA and XIIB in the Data Supplement**. *en face* immunofluorescent staining of abdominal aorta segments indicate Atg3 and p-eNOS^S1177^ protein expression are lower in ECs from iecAtg3KO vs. WT mice (**Figure 5C and 5D**, **Figure XIIC through XIID in the Data Supplement**). As a complementary approach, mRNA expression was assessed in ECs and M+A obtained from iliac arteries of both genotypes. Reduced *Atg3* gene expression was observed in pure iliac artery ECs (**Figure XIIE in the Data Supplement**) obtained from iecAtg3KO vs. WT mice. Substantiating the specificity of knockdown, mRNA expression of *Beclin1* was similar in ECs and M+A regardless of genotype (**Figure 5E**).

Since Atg3 compromise impaired shear-induced p-eNOS^S1177^ expression assessed *in vitro* and NO-mediated vasodilatory capacity examined *ex vivo,* we hypothesized that systemic blood pressure would be elevated in iecAtg3KO vs. WT mice and results from untethered 24 h biotelemetry support this (**Figure 5F and 5G**, **Figure XIIF through XIIG in the Data Supplement**). As would be predicted based on these results, medial thickness of the abdominal aorta was higher in iecAtg3KO vs. WT mice (**Figure XIII in the Data Supplement**), but indexes of perivascular fibrosis were not different between genotypes among heart, liver, or kidney (**Figure XIV in the Data Supplement**). These data indicate that knockdown of Atg3 specifically in ECs precipitates a hypertensive phenotype in adult mice.

### Impaired intraluminal flow-mediated vasodilation observed in femoral arteries from iecAtg3KO vs. WT mice is restored by 2-Me-ADP

Next we determined whether our results in HAECs with genetic autophagy compromise concerning shear-induced eNOS activation and NO generation can be translated to arteries from iecAtg3KO mice regarding intraluminal flow-mediated vasodilation. Animal and vessel characteristics are shown in SI Appendix, Table S5. Blunted vasodilation in response to three different flow rates was observed in arteries from iecAtg3KO vs. WT mice. Moreover, intraluminal incubation with 10^-3^ mol/L L-NMMA repressed flow-mediated vasodilation in arteries from WT but not iecAtg3KO mice, suggesting strongly that EC specific autophagy repression impairs NO-dependent vasodilation (**Figure 5H**). Importantly, incubation with 2-Me-ADP improved, but did not fully restore, vasodilatory capacity in arteries from iecAtg3KO mice (**Figure 5I**). Responses to sodium nitroprusside were similar in arteries from both genotypes indicating intact vascular smooth muscle function (**Figure VIE in the Data Supplement**). These findings illustrate that purinergic activation of autocrine signaling using 2-Me-ADP improves flow-induced vasodilation that is otherwise impaired in arteries from mice with EC specific depletion of Atg3 i.e., genetic loss of autophagy.

### P2Y_1_-R inhibition in adult mice mimics an aging vascular phenotype

Earlier we documented that shear-stress -induced p-eNOS^S1177^ and NO generation are prevented in BAECs treated with the P2Y_1_-R blocker MRS2179 ^7^. Using this loss of function approach, we sought to determine functional relevance. First we validated in HAECs that ADP-induced p-eNOS^S1177^ expression is negated by concurrent treatment with MRS2179 (**Figure 6A and 6B**) in the absence of cell death or apoptosis (SI Appendix, Figure S15). Supporting our hypothesis, intraluminal flow-mediated vasodilation displayed by arteries from adult mice in the presence of vehicle was markedly depressed during a second flow-response curve in the presence of MRS2179 (**Figure 6C**). Vascular smooth muscle function was not impacted (**Figure VF in the Data Supplement**). Of interest, limited vasodilatory capacity displayed by arteries from adult mice treated with MRS2179 was similar to that exhibited by vessels from old mice treated with vehicle or MRS2179.

**Figure 6.**
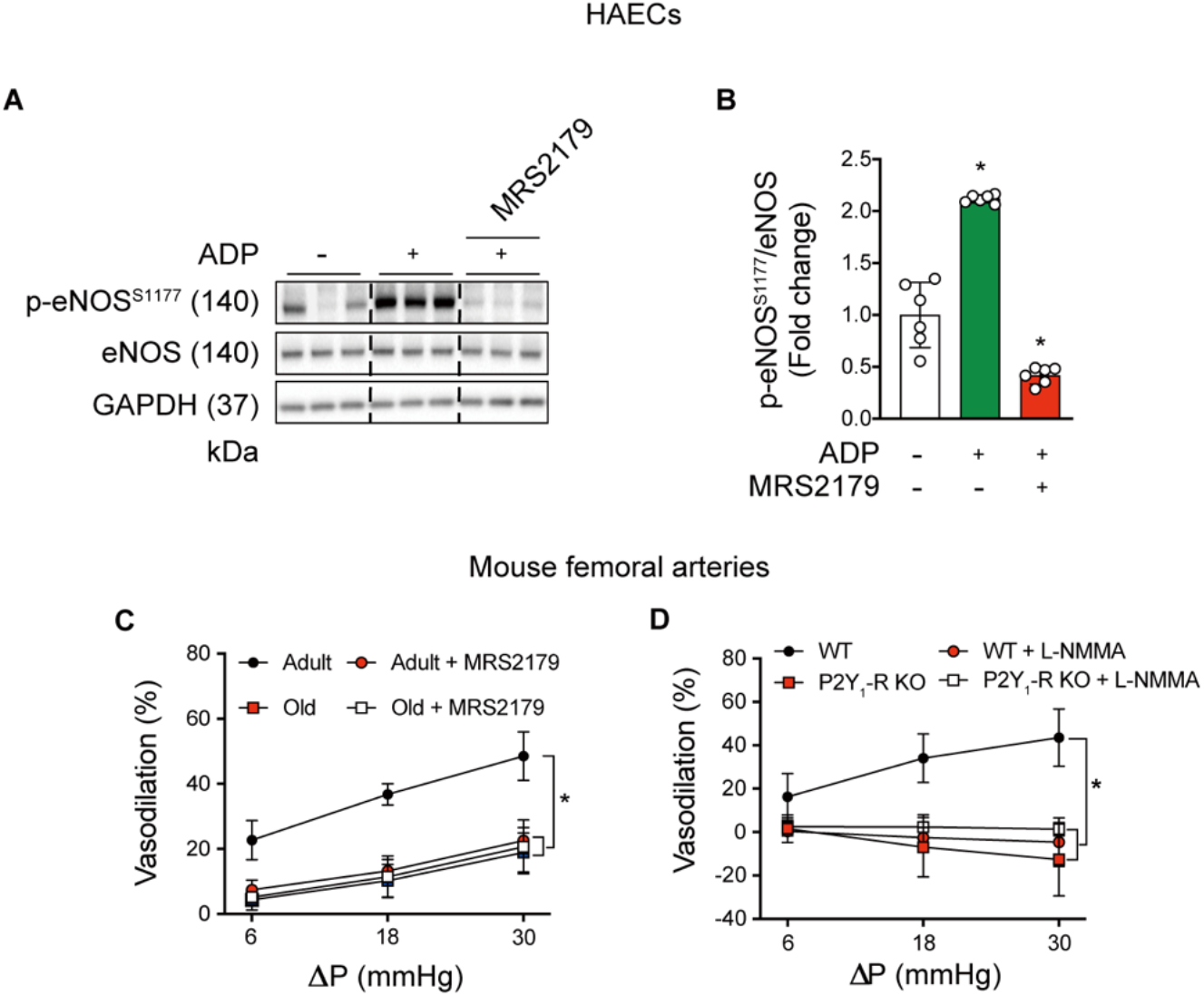
P2Y_1_-R inhibition in adult mice mimics an aging phenotype. Representative images (**A**) and mean densitometry (**B**) are shown. Adenosine diphosphate (ADP; 50 μmol/L) increased p-eNOS^S1177^ : eNOS protein expression in vehicle (PBS)-treated HAECs but not in HAECs that incubated with the P2Y_1_-R antagonist MRS2179 (5 μmol/L, 30 min). (**C**) Robust intraluminal flow-mediated vasodilation displayed by femoral arteries from adult mice is impaired by 30-min intraluminal incubation with 5 μmol/L MRS2179. Blunted vasodilatory capacity exhibited by arteries from older vs. adult mice is not repressed further by MRS2179. For **C**, n=3 mice per group, 1 artery per mouse. *p<0.05 vs Adult. (**D**) Intact intraluminal flow-mediated vasodilation displayed by arteries from WT but not P2Y_1_-R KO mice is nullified by 30-min incubation with L-NMMA. For **D**, n=4 mice per group, 1 artery per mouse. *p<0.05 vs WT. Statistical significance was assessed using a one-way ANOVA (**B**) and two-way repeated measures ANOVA (**C** and **D**).

Previously we reported that shear-stress induced p-eNOS^S1177^ and NO generation are prevented in BAECs transfected with P2Y_1_-R vs. scrambled siRNA ^7^. To determine functional relevance, we assessed intraluminal flow-mediated vasodilation in arteries from 2-month old male mice with germline depletion of P2Y_1_-Rs (i.e., P2Y_1_-R KO mice).^10^ Robust intraluminal flow-mediated vasodilation observed in femoral arteries from WT mice was absent in arteries from P2Y_1_-R KO animals, whereas vascular smooth muscle function was similar between groups (**Figure 6D**, **Figure VIG in the Data Supplement**). Notably, intraluminal flow mediated vasodilation displayed by arteries from WT mice was sensitive to NOS inhibition using L-NMMA, whereas vessels from P2Y_1_-R KO mice were resistant to this intervention. Animal and vessel characteristics are shown in **Table VI in the Data Supplement**.

When taken together, our findings provide solid evidence that physiological (aging), pharmacological (3-MA), and genetic (sgAtg3; iecAtg3KO mice) autophagy repression specifically in ECs compromises intraluminal flow-mediated vasodilation. Importantly, targeted activation of P2Y_1_-Rs via 2-Me-ADP in each context is sufficient to improve shear-stress -induced eNOS activation, NO generation, and / or arterial vasodilatory capacity. These findings translate our earlier results that 2-Me-ADP can reestablish shear-stress evoked NO generation in HAECs and BAECs that is otherwise prevented by pharmacological and genetic autophagy repression, to intraluminal flow-mediated vasodilation in arteries from mice with pharmacological, genetic, and aging-associated EC autophagy compromise.

## Discussion

The need is urgent to reveal new therapeutic targets for intervention in the context of aging-associated vascular disease. Here we report for the first time that : (i) older males with impaired FMD peak display repressed EC autophagy initiation and NO generation vs. adult males in response to elevated arterial shear rate associated with functional hyperemia; (ii) older mice with limited intraluminal flow-mediated arterial vasodilation exhibit blunted EC autophagy initiation and eNOS activation vs. adult mice in response to treadmill-running; (iii) inhibiting autophagy or disrupting P2Y_1_-Rs in the vasculature of adult mice mimics the age-associated defect concerning intraluminal flow-mediated vasodilation; and (iv) P2Y_1_-R activation rejuvenates intraluminal flow mediated vasodilation that is otherwise attenuated in arteries from mice after pharmacological, genetic, and aging associated EC autophagy compromise. These results provide solid evidence that age-associated arterial dysfunction concurrent with repressed EC autophagy can be reestablished by activating purinergic autocrine signaling.

### Older males with impaired vascular function display blunted shear-induced EC autophagy and eNOS activation in response to rhythmic handgrip exercise

While dysregulated autophagic activity is associated with aging-related pathologies including neurodegeneration, cancer, and immunosuppression, knowledge concerning the contribution from this process to aging-related cardiovascular complications in general^11–13^ and EC dysfunction in particular^2, 7, 8, 14, 15^, is evolving. From our search of the available literature, the influence of aging on arterial EC autophagy has been reported once in humans under basal conditions only.^5^ In that study, lower protein expression of beclin-1 together with p62 accumulation was observed in ECs obtained via brachial artery j-wire from older (61-71 y) vs. adult (20-31 y) subjects.^5^ We observed similar beclin-1 and p62 protein expression in radial artery ECs between groups at baseline (i.e., RHE-Pre), but LC3B, LC3B-bound puncta, and LC3B + LAMP2A colocalization were lower in ECs from older vs. adult participants. In general, both studies suggest several autophagy indexes are lower in ECs from older vs. adult subjects under basal conditions, but discrepant findings concerning beclin-1 and p62 are curious. EC biopsy procedures and immunofluorescent staining protocols were similar between reports, but whereas we used six male subjects per group, identified 75 radial artery ECs per endpoint, and normalized staining intensity to HAECs, LaRocca et al. examined five subjects (males and females) per group, identified 30 brachial artery ECs per endpoint, and controlled for staining intensity using human umbilical vein endothelial cells. Perhaps of greater importance is our novel finding that ECs from older subjects are unresponsive to physiological elevations of arterial shear rate concerning autophagy initiation and eNOS activation. Specifically, in addition to a lack of shear-induced p-eNOS^S1177^ and DAF fluorescence, ECs from older vs. adult volunteers were refractory to phagophore initiation (Beclin-1), autophagosome formation and maturation (Atg3 and LC3B puncta), autophagosome fusion with the lysosome (LC3B + LAMP2A colocalization), and lysosomal degradation of autophagosome contents (p62), in response to active hyperemia (**Figure 1 and 2**). Importantly, these results were obtained despite brachial artery blood flow velocity, absolute shear-rate, and the fold-increase in shear-rate above baseline being similar between adult and older subjects. Based on these data, the lack of eNOS activation and NO generation we observed previously in BAECs and HAECs exposed to shear stress after genetic and pharmacological autophagy compromise^7, 15^ appears to translate to primary arterial ECs from older humans with physiological repression of EC autophagy.

### Older mice with impaired vascular function display repressed EC autophagy and eNOS activation under basal conditions, and a failure to upregulate both processes in response to treadmill-running

In similar sized arteries, we confirm an age-associated repression of EC autophagy (iliac arteries) and intraluminal flow-mediated vasodilation (femoral arteries; **Figure 3**). Further, we report that aging mitigates the ability of arterial ECs to respond to functional hyperemia evoked by treadmill-running concerning EC autophagy activation. Specifically, 60-min of submaximal dynamic exercise elevated Atg3, LC3B, and p-eNOS^S1177^ in ECs from aorta of adult but not older mice (**Figure 4**). In light of these findings, we reasoned that a failure of flow/shear -induced EC autophagy initiation might limit eNOS activation and precipitate the impaired intraluminal flow-mediated vasodilation we observed in arteries from older vs. adult mice. In support of the contribution from NO to vasodilation, NOS inhibition abrogated vasodilatory capacity displayed by femoral arteries from adult mice with intact EC autophagy, but not older mice with compromised EC autophagy. However, because the circulating milieu associated with aging is complex, and factors other than depressed EC autophagy certainly contribute to endothelial dysfunction in this context, we tested and demonstrated that vascular autophagy inhibition *per se* (i.e., class III PI3K inhibition using 3-MA; **Figure 3F**), and EC specific autophagy depletion *per se* (i.e., iecAtg3KO mice; **Figure 5H and 5I**), are sufficient to mimic age-associated endothelial dysfunction. These data are the first to demonstrate that EC autophagy impairment is adequate to attenuate endothelial function. As such, age-associated reductions in arterial EC autophagy contribute importantly to limiting vasodilatory capacity exhibited by older rodents and humans.

### Age associated arterial dysfunction exhibited by mice with compromised EC autophagy is rejuvenated by promoting purinergic signaling

In a previous study we determined a molecular pathway that links repressed EC autophagy to impaired NO generation *in vitro.* Specifically, ECs with genetic disruption of autophagy exhibit diminished EC glycolysis and attenuated intra and extracellular ATP accumulation when exposed to shear-stress.^7^ Since studies using cultured ECs^16, 17^ and isolated cerebral arteries^18^ documented that ATP degradation products activate P2Y_1_-Rs and trigger NO generation, we evaluated this pathway further. Gain of function maneuvers that restored purinergic signaling (e.g., GLUT1 overexpression; 2-Me-ADP) rescued shear-induced p-eNOS^S1177^ and NO production in BAECs with impaired autophagy. Here we determined whether key results from BAECs studied *in vitro* can be translated to HAECs evaluated *in vitro* and arteries examined *ex vivo.* First we confirmed that 2-Me-ADP rescues shear stress-induced p-eNOS^S1177^ and NO generation in HAECs that is otherwise prevented by pharmacological (**Figure VII in the Data Supplement**) and genetic (**Figure X in the Data Supplement**) autophagy disruption. Demonstrating functional relevance of these *in vitro* studies, 2-Me-ADP improved intraluminal flow-mediated vasodilation that was otherwise attenuated in femoral arteries from aged mice with diminished basal EC autophagy and hyperemia-induced EC autophagy (**Figure 4**), and adult mice with EC specific depletion of autophagy (i.e., iecAtg3KO mice; **Figure 5**). If P2Y_1_-R activation preserves arterial function in a manner that is a consequence of defective autophagy, we predicted these responses should be resistant to pharmacological or genetic inhibition of autophagy. In support of this, otherwise blunted intraluminal flow-mediated vasodilation displayed by arteries from adult mice, older mice treated with 3-MA (**Figure 5A and B**), and adult iecAtg3KO mice (**Figure 5I**), is improved by concurrent treatment with 2-Me-ADP.

Loss of function approaches also were used in our previous study to test whether purinergic signaling serves as a link between repressed EC autophagy and impaired NO generation *in vitro.* For example, inhibiting glucose-transport via GLUT1 siRNA, blocking purinergic signaling via ectonucleotidase-mediated ATP/ADP degradation (e.g., apyrase), or inhibiting P2Y_1_-Rs using pharmacological (e.g., MRS2179) or genetic (e.g., P2Y_1_-R siRNA) procedures, prevented shear-induced p-eNOS^S1177^ and NO generation in ECs with intact autophagy.^7^ We sought to translate these results from ECs studied *in vitro* to arteries evaluated *ex vivo.* For example, if purinergic signaling to eNOS is an important component of intraluminal flow-mediated vasodilation in arteries from adult mice with intact autophagy, then disrupting P2Y_1_-R activation should precipitate a defect in the response. After validating that ADP-induced p-eNOS^S1177^ is negated by concurrent treatment with MRS2179 in HAECs, we demonstrated that robust intraluminal flow-mediated vasodilation displayed by arteries from adult mice could be attenuated by luminal incubation with this P2Y_1_-R blocker, to values not different from those exhibited by arteries from older mice (**Figure 6**). These results were substantiated using arteries from mice with germline P2Y_1_-R deficiency.

### Conclusions

Evidence from our laboratory and others indicate aging is associated with repressed vascular autophagy and impaired NO-mediated arterial function, but a confirmed link has not been established. eNOS is regulated at multiple levels,^19^ including via nucleotide activation of purinergic receptors.^16, 20, 21^ Our earlier findings *in vitro* indicate that genetic repression of autophagy precipitates defects in purinergic signaling to eNOS that can be rescued. Here we provide novel evidence that dysfunction displayed by aged arteries with repressed EC autophagy can be reestablished by pharmacological promotion of purinergic autocrine signaling. Further research concerning the G protein-coupled P2Y family of receptors is warranted in an effort to explore new therapeutic treatment options for aging-related endothelial dysfunction that is secondary to or associated with defective EC autophagy.

## Acknowledgments

JMC, SKP, JDT, LPB, SB, JDS contributed to the conception and design of the study. JMC, SKP, OSK, DTLS, JC, MT, JDT, JDS performed experiments and data acquisition. DM, AN, AB collected endothelial cells from human subjects. FW and TXY performed biotelemetry experiments. SPK contributed reagents. JMC, SKP, JDS wrote the manuscript. All authors provided important intellectual contributions.

