## Supplementary material for "Activating P2Y_1_ receptors improves function in arteries with repressed autophagy": Supplmental Tables, Figures, Video, and Methods: Sup Manuscript (ATVB)-BIORxiv.pdf

[1] Department of Nutrition and Integrative Physiology, University of Utah, Salt Lake City, UT, USA
[2] Division of Endocrinology, Metabolism and Diabetes, and Program in Molecular Medicine University of Utah School of Medicine, Salt Lake City, UT, USA
[3] Department of Kinesiology, University of Connecticut, Storrs, CT, USA
[4] Department of Orthopedic Surgery & Center on Aging, University of Connecticut School of Medicine, Storrs, CT, USA
[5] Geriatric Research, Education, and Clinical Center, George E. Whalen Veterans Affairs Medical Center, Salt Lake City, UT, USA
[6] Department of Anesthesiology, University of Utah, Salt Lake City, UT, USA
[7] Sol Sherry Thrombosis Research Center, Lewis Katz School of Medicine, Temple University, Philadelphia, PA, USA
[8] Department of Internal Medicine, Division of Geriatrics, University of Utah, Salt Lake City, UT, USA
[9] Nephrology Research, George E. Whalen VA Medical Center, Salt Lake City, UT, USA
[10] Department of Internal Medicine, Division of Nephrology, University of Utah, Salt Lake City, UT, USA
* Equal contribution

#### Supplemental Methods

##### *Human studies*

Protocol approval and written informed consent were obtained according to the University of Utah and Salt Lake City Department of Veterans Affairs Medical Center Institutional Review Boards. Adult and older male subjects that were sedentary to recreationally active, nonobese, nonsmokers, free of diagnosed cardiovascular or metabolic complications, and were not taking any medications, participated in the study. Subjects reported to the laboratory on two occasions, separated by less than 7-days, having abstained from caffeine, alcohol, and exercise for 24 h.

##### *Flow mediated vasodilation and maximal handgrip workload test*

During visit 1 a flow-mediated vasodilation (FMD) test was completed using procedures we have described.[1] After a 20-min rest period, a maximal handgrip workload test was completed to estimate an intensity that would result in a ~ 3-fold elevation of arterial shear rate during rhythmic handgrip exercise (RHE). Subjects lay supine with their right arm extended perpendicular to their body, while their right hand grasped a handle attached to a pulley connected to a bucket containing 0.9 kg (**Video I in the Data Supplement**). When the subject squeezed and released the handle, once every two sec, a weight-containing bucket elevated and descended 3.5 cm, respectively. Additional weight (0.9 kg) was placed in the bucket each minute (i.e., every 30 contractions) until the subject could no longer perform the task or began recruiting accessory muscles to complete the maneuver.[1]

##### *EC collection and processing*

Visit 2 occurred within 7 days of visit 1. A 20 g, 5 cm catheter (Arrow, RA-04020) was inserted into the radial artery of the right arm under aseptic conditions. Immediately after catheter insertion, a blood pressure cuff was placed to occlude the brachial artery and thus radial artery blood flow. A flexible 0.025 inch mesh 3-mm guide wire with a J-shaped tip (i.e., j-wire; GuideRight) was advanced ~3-4 cm beyond the termination of the radial artery catheter and retracted.[1-3] The distal portion of the j-wire was placed into dissociation buffer [0.5% BSA, 2mmol/L EDTA, 18 U/ml heparin in PBS (pH 7.4)]. This represents the RHE-Pre collection. Subjects then completed the 60-min RHE protocol (see below). Immediately upon the cessation of RHE, EC's were obtained by advancing the j-wires 4-6 cm beyond the end of the catheter. This represents the RHE-Post collection. Each j-wire was advanced different distances beyond the end of the arterial catheter at RHE-Pre and RHE-Post collections so that cells could be obtained from different areas. Within 30-min of the RHE-Pre and RHE-Post collection, the admixture of cell types was dislodged from each j-wire by washing with dissociation buffer. Cells were recovered and treated with an erythrocyte lysing kit. Next, cells were resuspended in EBM-2 (Lonza Inc), and applied evenly to chamber slides pre-treated with poly-L-lysine. Gentle centrifugation (450 rpm / ~28g, 10 s with deceleration set to 0, Thermo Fisher Scientific), in a microplate swinging-bucket rotor, facilitated adherence of the cells to the slide surface. A list of all reagents and their sources is listed in the *Major Resources Table.*

##### *Assessment of nitric oxide (NO) and superoxide anion ($O_2^{\bullet-}$) generation*

A cohort of cells was stained on the day of collection without fixation to estimate NO and $O_2^{\bullet-}$ generation.[1,4] Cells were incubated for 30 minutes at 37°C with either 5 μmol/L 4-amino-5-methylamino-2',7'-difluorofluorescein diacetate (DAF-FM Diacetate), to estimate NO generation, or 10 μmol/L dihydroethidium (DHE), to estimate $O_2^{\bullet-}$ production. After incubation, all cells were co-stained with VE-cadherin and 4',6-diamidino-2-phenylindole (DAPI) to identify ECs and nuclei, respectively. In preliminary studies using HAECs pyocyanin-induced elevations in $O_2^{\bullet-}$ generation were inhibited using N-acetyl-L-cysteine, and insulin-induced stimulation of NO generation was ameliorated using L-NMMA (data not shown).

***Assessment of protein expression : immunofluorescence.***
A cohort of cells was fixed with 4% paraformaldehyde (PFA; Sigma-Aldrich) on the day of collection, and frozen at -80°C until staining for protein expression using immunofluorescent antibodies.[1, 4] Commercially available HAECs (P4-6, Lonza Inc) were treated in an identical manner to serve as intensity controls. On the day of staining fixed cells were thawed and permeabilized in 0.1% Triton X-100 and nonspecific binding sites were blocked in 0.5% BSA. Cells were incubated for 1 h at room temperature with primary antibodies to two of the following targets: p-eNOS$^{S1177}$; eNOS; Beclin-1; Atg3; LC3B; p62; and LAMP2A. Cells were simultaneously stained with VE-cadherin to identify ECs. Slides next incubated with conjugated secondary antibody (i.e., Alexa Fluor 488, Alexa Fluor 555, and Alexa Fluor 647) for 30 minutes at room temperature. Finally, slides were mounted with ProLong Diamond Antifade mount with DAPI for nuclear identification. Antibody source, catalogue numbers, and dilutions are documented in the the *Major Resources Table*, Table S8. All slides were imaged with a confocal microscope (A1R, Nikon) at x 60 magnification. Images were captured at the same exposure time and corrected for background fluorescence. Seventy-five ECs were identified on each slide by positive co-staining for VE-cadherin and DAPI. Images were captured / analyzed with NIS element AR software. Fluorescence intensity was measured using ImageJ (NIH). For protein expression, data are expressed as the ratio of RHE-Pre or RHE-Post staining intensity divided by HAEC staining intensity x 100 to control for differences between staining sessions.[1] For NO and $O_2^{\bullet-}$ data are expressed as arbitrary units (AU).

***RHE protocol and hemodynamics.***
RHE was performed during visit 2 to increase radial artery shear rate using procedures that we have described.[1] Thirty-min after the RHE-Pre EC collection, subjects squeezed and released a handle that elevated a bucket containing 5-10% of the weight achieved during their maximal handgrip workload test completed during visit 1. The duty cycle was 1 : 2 and subjects contracted to the sound of a metronome (**Video I in the Data Supplement**). The weight was designed to provide a workload that required ~ 3-fold elevations in radial artery shear rate that could be sustained for 60-min. The elevation of radial artery shear rate was estimated throughout the 60-min RHE protocol by directly measuring brachial artery (BA) shear rate (see below) over 6 cardiac cycles at 10 min intervals. Rating of perceived exertion (RPE) was assessed according to the modified 10 point Borg scale i.e., mild (0-3), moderate (4-7), or difficult (8-10). Using BA diameter and BA blood velocity ($V_{mean}$) collected using Doppler ultrasound, BA blood flow [($V_{mean}\pi$ (vessel diameter / 2)$^2$ x 60] and shear rate ($8V_{mean}$/BA diameter) were calculated.[1] Heart rate was monitored using a standard 3-lead ECG, stroke volume and cardiac output by photoplethysmography, and mean arterial pressure with an automated sphygmomanometer throughout the protocol.[1] Hemodynamic variables before and during RHE were averaged over 6 cardiac cycles at 10-min intervals.

**Human aortic endothelial cells (HAECs)**

HAECs (Lonza Inc) were maintained in endothelial basal medium 2 (EBM2; Lonza Inc) containing supplements (EGM-2 SingleQuots, Lonza Inc) in a 5% $CO_2$ atmosphere at 37°C. HAECs were passaged at 70-80% confluency and passages 4-6 were used for the respective experiments.[1, 5]

First we confirmed the ability of the class III phosphatidylinositol 3-kinase (PI3K) inhibitor 3-methyladenine (3-MA) to inhibit autophagy in the context of our experimental conditions. HAECs were exposed to 0 or 20 dyn/cm$^2$ for 45-min, in the absence and presence of 5 mmol/L 3-MA. This intensity of shear stress is relevant physiologically because arterial shear forces range from ~ 5 dyn/cm$^2$ in descending aorta to ~ 55 dyn/cm$^2$ in arterioles.[4, 6, 7] Shear stress was estimated as: shear stress = $\alpha \sqrt{(p*\eta(2\pi f)^3)}$, where α is the orbital radius of rotation (2.5 cm), p is the medium density (1.01 g/ml), η is the viscosity of the medium [0.01 poise measured with a rheometer (TA Instruments Inc, model AR550)], and f is the frequency of rotation (2.92 rotations/sec).[4, 6]

Second we substantiated that shear-induced p-eNOS$^{S1177}$ and NO generation are inhibited by 3-MA in a manner that can be restored by the purinergic 2Y1 receptor agonist 2-methylthio-ADP (2-Me-ADP). HAECs incubated in the absence and presence of: (i) 5 mmol/L 3-MA; (ii) 100 µmol/L 2-Me-ADP; or (iii) 5 mmol/L 3-MA + 100 µmol/L 2-Me-ADP.[4]

Third we confirmed the ability of the $P2Y_1$-R antagonist MRS2179 to inhibit ADP-induced p-eNOS$^{S1177}$. To do so, HAECs were treated with 50 µmol/L ADP for 1 min in the absence or presence of 5 µmol/L MRS2179. Immunoblotting was performed as described below.

Fourth, in addition to pharmacological repression of EC autophagy using 3-MA, we used CRISPR-Cas9 to deplete Atg3 in HAECs. CRISPR MIT software (http://crispr.mit.edu/) was used to design a highly efficient and specific target sequence 5'-ACAAACGTGGCGAATAT-3' which was ordered from Synthego. Twenty pmol/L of Cas9 2NLS (Synthego) was mixed with 60 pmol/L of synthetic sgRNA dissolved in nuclease-free 1X TE buffer to synthesize the Cas9 RNP complex. The mixture was incubated for 10 min to assemble the RNP complex. HAECs (1x10$^5$) were harvested, washed with PBS, and the pellet was resuspended in 10 µl resuspension buffer R (Thermo Fisher Scientific). Equal volumes of cell suspension buffer R and RNP complex were gently mixed, and the solution was aspirated into the Neon tip (Thermo Fisher Scientific). After transfection (Neon Transfection System; Thermo Fisher Scientific; 1400 V, 20 ms, 2 pulses), cells were transferred to a 6-well plate. Shear-induced indexes of autophagy, p-eNOS$^{S1177}$, and NO generation were measured in wild type (WT) HAECs and sgAtg3 HAECs in the absence and presence of 2-Me-ADP.[4] After protocols 1-4 described above, immunoblotting procedures were performed as described below.

***NO generation***

NO generation was estimated in HAECs in the absence and presence of: (i) 5 mmol/L 3-MA; (ii) 100 µmol/L 2-Me-ADP; or (iii) 5 mmol/L 3-MA + 100 µmol/L 2-Me-ADP, without (0 dyn/cm$^2$) or with (20 dyn/cm$^2$) shear stress for 45-min.[1, 4, 6] HAECs incubated with 5 µmol/L DAF-FM Diacetate. After the respective treatments, HAECs were co-stained with DAPI to identify nuclei. HAECs were imaged with an automated wide-field fluorescence microscope (Nikon) at x10 magnification, and images were captured using an ultra-high sensitivity Andor Clara CCD camera. Fluorescence intensity was quantified with Image J software (NIH).

***Cell viability***

In response to all treatments, HAECs were cultured as described and seeded on a 96-well plate in EGM. HAECs were treated with the respective vehicle or pharmacological agent. To determine cytotoxicity, cells incubated with 10 µl of PrestoBlue reagent for 30 min. PrestoBlue is modified by the reducing environment of the viable cell, turns red, and becomes highly fluorescent. Absorbance was read at 570 nm using a spectrophotometer, and values were normalized to the 600 nm values for the experimental wells. The FITC Annexin V/Dead Cell Apoptosis Kit was used to assess apoptosis using guidelines provided by the manufacturer. After the respective treatments, ECs were detached and suspended in 1 X binding buffer, and treated with Alexa Fluor 488 Annexin V and propidium iodide (PI). In apoptotic cells, phosphatidylserine (PS) is translocated from the inner to the outer leaflet of the plasma membrane, thus exposing PS to the external cellular environment. Annexin V labeled with FITC has a high affinity for PS exposed on the outer leaflet. PI is impermeant to live cells and apoptotic cells, but stains dead cells with red fluorescence, binding tightly to the nucleic acids in the cell. After staining, apoptotic cells show green fluorescence, dead cells show red and green fluorescence, and live cells show little or no fluorescence. Cell populations were distinguished using a flow cytometer (Becton Dickinson) with the 488 nm line of an argon- ion laser for excitation into four quadrants : necrotic cells (Q1), late apoptotic cells (Q2), early apoptotic cells (Q3), and viable cells (Q4).[4, 5]

***Animal studies***
Procedures involving mice were approved by the Institutional Animal Care and Use Committee at the University of Utah.

*Older vs. adult mice.* Male C57BL/6 mice were obtained from the Jackson Laboratories at 5-6 months of age, or the National Institute on Aging rodent colony at 21 months of age, and housed in AALAC accredited facilities at the University of Utah. Animals were maintained on a 12 : 12 h light : dark cycle in a temperature controlled environment (22-23°C), and were provided standard rodent chow and water ad libitum. Separate cohorts of adult (7 months) and old (24 months) mice were used to assess: (i) EC and media + adventitia (M+A) mRNA expression and arterial (aorta, iliac, femoral) protein expression of autophagy-related genes via qRT-PCR and immunoblotting, respectively; (ii) intraluminal flow-mediated vasodilation of femoral arteries examined *ex vivo* via myography; and (iii) indexes of autophagy and eNOS activation in arterial ECs via *en face* staining. Male, 23-24 month old C57BL/6 mice are a well-accepted pre-clinical model of age-associated vascular dysfunction.

***Mice with inducible depletion of Atg3 specifically in ECs***
Atg3 mediates lipidation of Atg8/LC3I to form mature LC3II and is required for the formation of autophagosomes and the activation of autophagy. Flox-Atg3 mice were developed in collaboration with the University of Utah Transgenic and Knockout Mouse Core Facility as we described [4]. Atg3$^{flox/flox}$ mice were crossed with Cdh5-Cre$^{ERT2}$ mice that were kindly provided by Dr. Ralf Adams [8]. Atg3$^{flox/flox}$ and Cdh5-Cre$^{ERT2}$ mice were on a C57BL/6 background. Atg3$^{flox/flox}$ / Cdh5-Cre$^{ERT2}$ (iecAtg3KO) and Atg3$^{flox/flox}$ littermates (WT) were used in this study. Floxed Atg3 and Cdh5-Cre were confirmed by PCR analysis of genomic DNA. The sequences of primers are listed in **Table VII in the Data Supplement**. To activate the Cdh5-Cre in 4-month old mice, 4 mg tamoxifen was administered via oral gavage to iecAtg3KO and WT mice for 4 consecutive days. Body weights were not significantly different before or after the 4-day dosing regimen. Separate cohorts of WT and iecAtg3KO mice were used to assess: (i) EC and M+A mRNA

expression of autophagy-related genes via qRT-PCR; (ii) intraluminal flow-mediated vasodilation of femoral arteries examined *ex vivo* via myography; and (iii) indexes of autophagy and eNOS activation in arterial ECs via *en face* staining.

***Mice with germline depletion of purinergic 2Y1 receptors***

Two-month old $P2Y_1$-R knockout mice ($P2Y_1$-R KO mice) and their wild type (WT) littermates that have been characterized previously were used.[9] Femoral arteries from these animals were used to assess intraluminal flow-mediated vasodilation examined *ex vivo* via myography.[5, 10, 11]

***Tissue collection***

A blood sample from conscious, random-fed mice was used to assess glucose. Next, mice were anesthetized using 2-5% inhaled isoflurane combined with 100% oxygen. When a stable plane of anesthesia was attained, the chest was opened using aseptic procedures, the heart was exposed, and a small incision was made in the right atrium. A 23-gauge (g) needle attached to an insulin syringe was used to slowly deliver 1 ml of phosphate buffered saline containing 1000 U / ml heparin into the left ventricle. Next, the heart was excised, and the carotid arteries, entire aorta, iliac arteries, and femoral arteries were cleaned of adherent tissue while bathed in iced physiological saline solution (PSS) containing (mmol/L): 145.0 NaCl, 4.7 KCl, 2.0 $CaCl_2$, 1.17 $MgSO_4$, 5.0 glucose, 2.0 pyruvate, 0.02 EDTA, 3.0 MOPS buffer, 10 g/ L of BSA at pH 7.4, and protease and phosphatase inhibitors.[4, 5, 10-12]

***mRNA expression***

At the time of tissue collection for experimentation, carotid and iliac arteries were obtained from adult and old mice, and iecAtg3KO and WT mice. Both segments of each vessel type were excised and perfused with 200 μl QIAzol lysis reagent (QIAGEN) using a blunt-end 27 g needle.[4] The vessel effluent containing the intimal fraction (i.e., ECs) was collected. The remainder of the vessel (media + adventitia; M+A) was placed in a separate tube. To confirm purity of each fraction, mRNA expression of *Pecam1* and *α-Sma* in ECs and media + adventitia was assessed using qRT-PCR and SYBR green fluorescence (Qiagen, #204143), and cycle threshold values were normalized to *18S*.[4] *18S* was chosen based on preliminary studies wherein its stability over time i.e., 7 and 24 months, was superior relative to *B2m*, *Hprt*, *Actb*, and *Gapdh*, using geNorm software analysis (Biogazelle).[13] The average expression stability value of *18S* was lower than 0.5. A list of all primers used in this study is shown in **Table VIII in the Data Supplement**.

***En face immunofluorescence***

We used *en face* staining of ECs in segments of aorta from : (i) adult and old mice that did (EX) or did not (No-EX) complete 60-min of treadmill-running at an intensity designed to evoke 70% of their maximal workload capacity (see below); and (ii) adult WT and iecAtg3KO mice 14 days after the final tamoxifen administration. At the time of tissue collection, segments of aorta were obtained, permeabilized using 0.1% Triton X-100, blocked using 10% BSA for 30 min, and stained using fluorescent antibodies specific to Atg3, LC3B, p-$eNOS^{S1177}$, VE-Cadherin and DAPI. 5 images per aorta segment per mouse were acquired using confocal microscopy (A1R, Nikon) at x 60 magnification. Images were analyzed using NIS element AR software. The intensity of staining for the respective protein in cells co-staining with VE-Cadherin and DAPI was normalized to cell numbers. Twenty cells from each image were quantified.

***Acute exercise***

Cohorts of adult and old mice used for *en face* staining of ECs before and after acute exercise were familiarized with walking/running on a motorized treadmill (Columbus Instruments) for two consecutive days. On day 3, a workload capacity evaluation test was completed on each mouse. Total workload was calculated as [body weight (kg) x total running time (min) x final running speed (m/min) x treadmill grade (25%)]. Electric shocks were not used. Mice were encouraged to run by tapping their rear using test tube cleaning brushes. At least 24 h later mice were separated into a group that did (EX) or did not (No EX) complete 60-min treadmill-running at a speed and grade designed to achieve 70% of their maximal workload. Mice that did not complete acute exercise remained in their home cage beside the treadmill so that they were exposed to sounds / vibrations associated with treadmill-running.

***Protein expression***

In HAECs that completed treatments described earlier in *Cell studies*, and in aorta, iliac, and femoral homogenates from adult and old mice, protein expression was measured as described.[4, 5, 10-12] Briefly, HAECs or arterial homogenates were lysed in RIPA buffer containing protease and phosphatase inhibitors and EDTA. Protein (35 µg) was resolved in 6X sample buffer and loaded into 4-20% SDS-PAGE gels (Bio-Rad, #5671094). Next, the proteins were transferred to a nitrocelluose membrane using a semi-dry transfer device (iBlot2, Thermo Fisher Scientific). The membrane was blocked using 5% skim milk for 1 hour at room temperature, and incubated with the following primary antibodies; p-eNOS$^{S1177}$, eNOS, p62, LC3B, GAPDH and secondary antibodies; Goat anti-Rabbit, Goat anti-Mouse. Protein bands were detected by chemiluminescence and subsequently quantified by ImageJ (NIH).

***Intraluminal flow-mediated vasodilation***

Segments of femoral artery were immersed in 4°C PSS. Each end of the vessel was cannulated using a micropipette tip with the aid of a dissecting microscope (SZX10; Olympus). After the temperature of the bathing medium was increased over 30-min to 37°C, arteries equilibrated for 1 h, followed by 10 mmHg increases in intraluminal pressure every 5-min to 60 mmHg. Five responses were observed in each femoral artery. First, a cumulative concentration response curve to potassium chloride (KCl, 20 - 100 mmol/L) was obtained to measure non-receptor mediated vasoconstriction. Second, to measure receptor-mediated vasoconstriction, a cumulative concentration response curve to phenylephrine (PE, $10^{-8}$ M-$10^{-5}$ mol/L) was obtained, and the dose of PE required to evoke 50% of maximal PE-evoked constriction was calculated. Third, after intraluminal incubation with the respective vehicle, and upon 50% of maximal PE-induced vasoconstriction, intraluminal flow was initiated by increasing the inflow pressure (e.g., P1) and decreasing the outflow pressure (e.g., P2) to create a pressure gradient (e.g., ΔP) across the vessel. For example, from isobaric conditions wherein P1=60 mmHg =P2, a ΔP of 6 mmHg is created by increasing P1 to 63 mmHg and decreasing P2 to 57 mmHg. Percent vasodilation was measured in response to ΔP's of 6 mmHg, 18 mmHg, and 30 mmHg x 3-min each. Fourth, after arteries equilibrated in an isobaric environment (60 mmHg) for 30-min in the presence of intraluminal : (i) NOS inhibition using L-NMMA; (ii) autophagy inhibition using 3-MA (5 mmol/L); (iii) $P2Y_1$-R blockade using MRS2179 (5 µmol/L); (iv) $P2Y_1$-R activation using 2-Me-ADP (100 µmol/L); or (v) a combination of 3-MA and 2-Me-ADP, a second intraluminal flow-mediated vasodilation curve was completed after stable PE-induced precontraction. Fifth, after arteries equilibrated at 60 mmHg for 30-min, and upon PE-induced precontraction, a concentration-response curve to the endothelium-independent vasodilator sodium nitroprusside (SNP, $10^{-9}$ - $10^{-4}$ mol/L) was completed to estimate vascular smooth muscle function.

For all interventions, percent vasodilation was calculated as (DT-Dp) / (Di-Dp) × 100. Where DT is the recorded diameter at a given time point (i.e. diameter response to flow or SNP), Dp is the diameter recorded after the addition of the vasoactive agent (i.e. pre-constriction diameter), and Di is the diameter recorded immediately before the addition of the vasoactive agent (initial diameter). Percent vasoconstriction (% of baseline) was calculated as Dp / Di x 100.[5, 10, 11]

***Blood pressure***
WT and iecAtg3KO mice were anesthetized using 2% isoflurane combined with 100% oxygen. Using aseptic techniques, the neck area was shaved, disinfected, and a catheter was inserted into the left common carotid artery. A transmitter probe (Data Sciences International) attached to the catheter was secured in the abdomen in a manner that allowed ambulation with minimal hindrance. Thirty second averages of blood pressure (systolic, diastolic, mean) and heart rate (HR) were recorded every 15-min, for three 24-hour periods, starting 6 days post-surgery. Three 24-hour periods (i.e., days 7-9 post-surgery) were averaged into one 24-hour period and used for data analysis.[5, 10, 11]

***Histology***
Segments of aorta from adult, old, WT, and iecAtg3KO animals, together with heart, liver, and kidney from WT and iecAtg3KO mice, were obtained and placed immediately in 4% PFA. Next, tissues were dehydrated, embedded in paraffin, sectioned (3 µm thick), mounted on glass slides, and stained with Masson's Trichrome at the University of Utah Histology Core Facility. Perivascular fibrosis was assessed in organs from WT and iecAtg3KO mice using 30 fields of view. Intima and media thickness was assessed in segments of aorta from all groups using 9 fields of view. All images were acquired via Olympus IX71 at X10 magnification, and the data were analyzed using the CellSens Dimension Program.

***Statistical analyses***
Data are presented as mean ± standard deviation of the mean. Significance was accepted when $p < 0.05$. A test to determine normality of distribution for each data set was performed by GraphPad Prism software version 9. For comparison between two groups, unpaired or paired t-tests were performed as appropriate. For comparison among three or more mean values, if the data were distributed normally, a one-way ANOVA was performed to determine whether significant differences exist among groups. If significance was obtained, a Tukey's post hoc test was used to identify the location of the differences. If the data were not distributed normally, a Kruskall-Wallis ANOVA was performed. If significance was obtained, a Dunn's post hoc test was used to identify the location of the differences. Two-way ANOVA was used to determine significance concerning: (i) RHE-Pre and RHE-Post between adult and old subjects; and (ii) intraluminal flow-mediated vasodilation in arteries. A two-way repeated measured ANOVA was used to determine significance concerning hemodynamic variables during the rhythmic handgrip exercise protocol. Note that information concerning statistical tests are included in each figure and table legend.

#### Supplemental Figures and Figure Legends

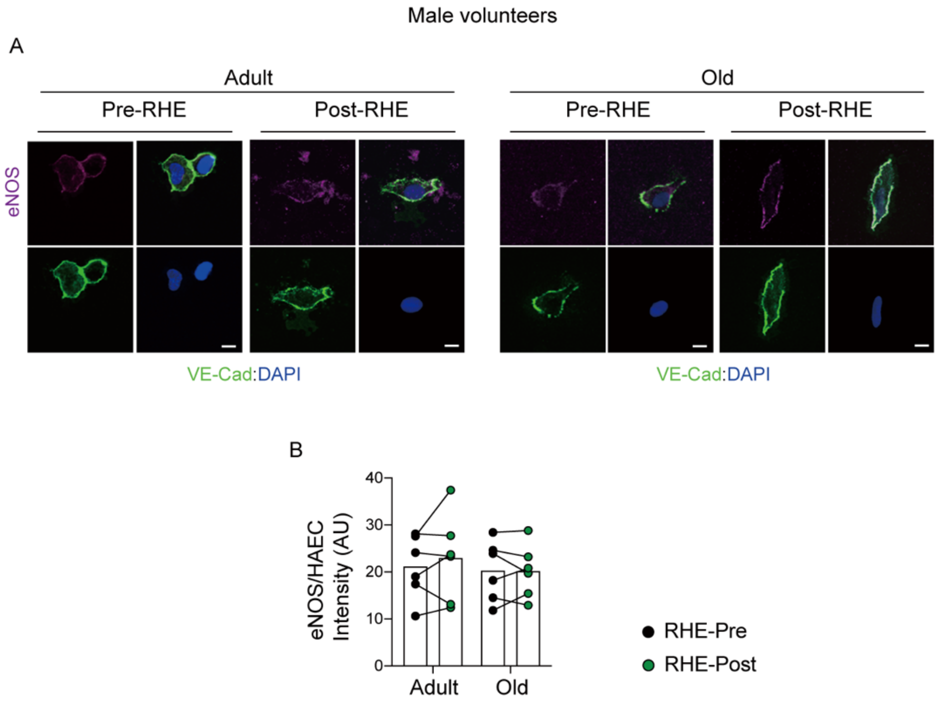

**Supplemental Figure I. eNOS protein expression is similar in adult and older subjects.** Representative images (**A**) and mean immunofluorescence staining (**B**) is shown. Comparing RHE-Pre to RHE-Pre between groups i.e., the influence of aging, eNOS intensity is similar in ECs from older vs. adult participants (i.e., histogram 1 vs. 3). Comparing RHE-Pre to RHE-Post i.e., the influence of elevated arterial shear rate, eNOS is similar in ECs from adult (bar 1 vs. 2) and older (bar 3 vs. 4) subjects. Seventy-five ECs from each time point and each subject were measured, and the staining intensity was normalized to values obtained from commercial HAECs using identical conditions. Scale bar represents 10 μm. Magnification=60 X. Statistical significance was assessed using a two-way repeated measures ANOVA.

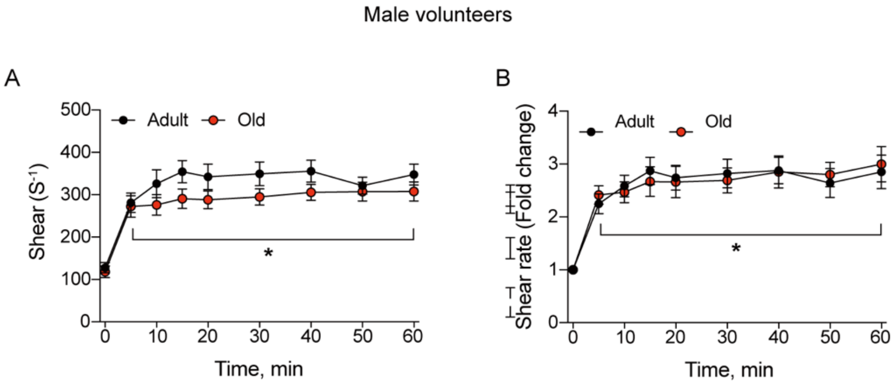

**Supplemental Figure II. Rhythmic handgrip exercise increased arterial shear rate similarly in adult and older subjects.** Absolute shear rate (**A**) and the fold-increase in shear-rate from 0-min (**B**) evoked by rhythmic handgrip exercise was not different between groups. For **A** and **B**, n=6 per group. Statistical significance was assessed using a two-way repeated measures ANOVA.

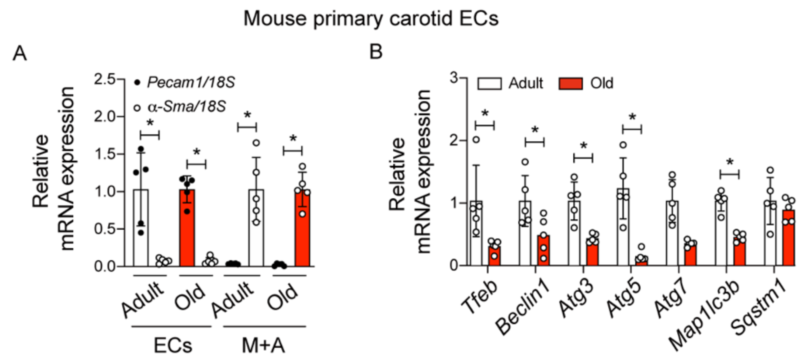

**Supplemental Figure III. EC autophagy is repressed in carotid arteries from old vs. adult mice.** ECs and media + adventitia (M+A) were obtained from carotid arteries of adult and old mice using procedures described in the text. Purity of the EC and M+A fraction was verified by quantifying mRNA expression of platelet/endothelial cell adhesion molecule 1 (*Pecam1*) and alpha-smooth muscle actin (*α-Sma*), respectively. *Pecam1* mRNA was highly expressed in ECs, but not in M+A, whereas *α-Sma* mRNA was highly expressed in M+A, but not in the EC fraction (**A**) Using the EC fraction only from carotid arteries, we demonstrate that mRNA expression of transcriptional factor EB *(Tfeb*), *Beclin1, Atg3, Atg5,* and *Map1lc3b* is lower in older vs. adult mice (**B**).

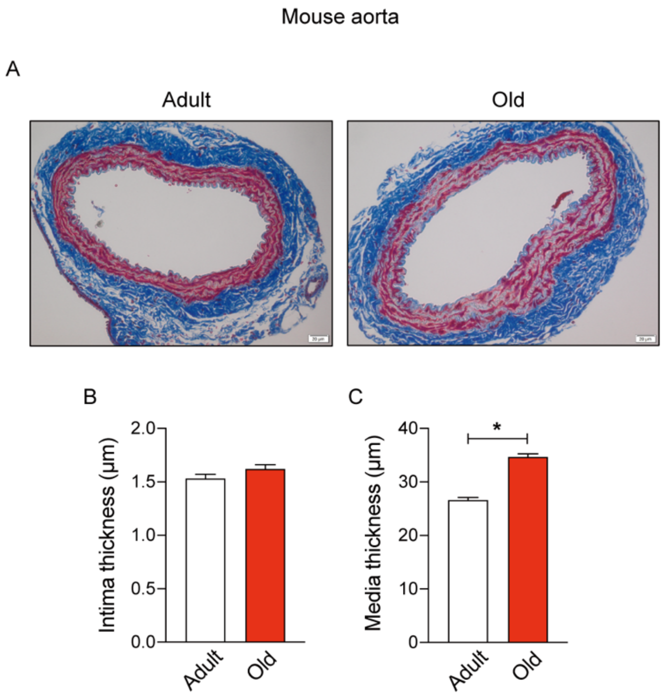

**Supplemental Figure IV. Media thickness is greater in aorta from old vs. adult mice.** A representative image of trichrome staining (**A**) and mean data concerning: (**B**) intima thickness; and (**C**) media thickness. Scale bar : 20 µm. Magnification : 10X. For **B** and **C**, n=5 mice, one segment of aorta per mouse, obtained just proximal to the bifurcation of the iliac arteries. *p<0.05 vs adult. Statistical significance was assessed using an unpaired t-test.

Mouse femoral arteries

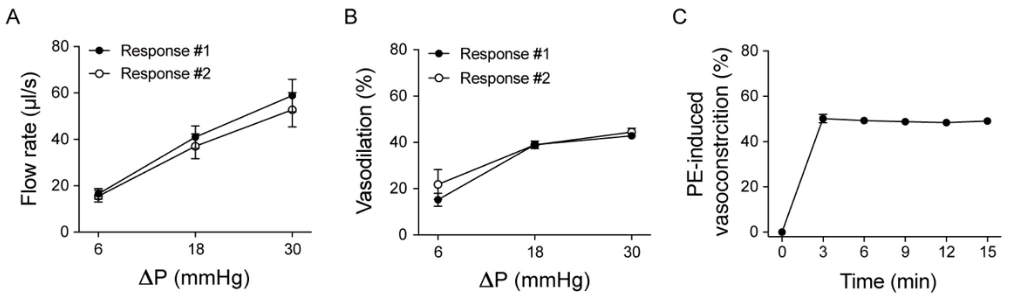

**Supplemental Figure V. Control studies for vascular function experiments.** (**A**) Similar flow rates are evoked by ΔPs of 6, 18, and 30 mmHg when responses are separated by 30-min. (**B**) Tachyphylaxis is avoided if intraluminal flow-mediated vasodilation responses are separated by 30-min. (**C**) Phenylephrine-evoked vasocontraction is stable for a duration (e.g., 15-min) sufficient to complete each of the vasodilation experiments that are described herein. These control experiments were completed using 3 adult mice, 2 femoral arteries per mouse. Statistical significance was assessed using a two-way repeated measures ANOVA.

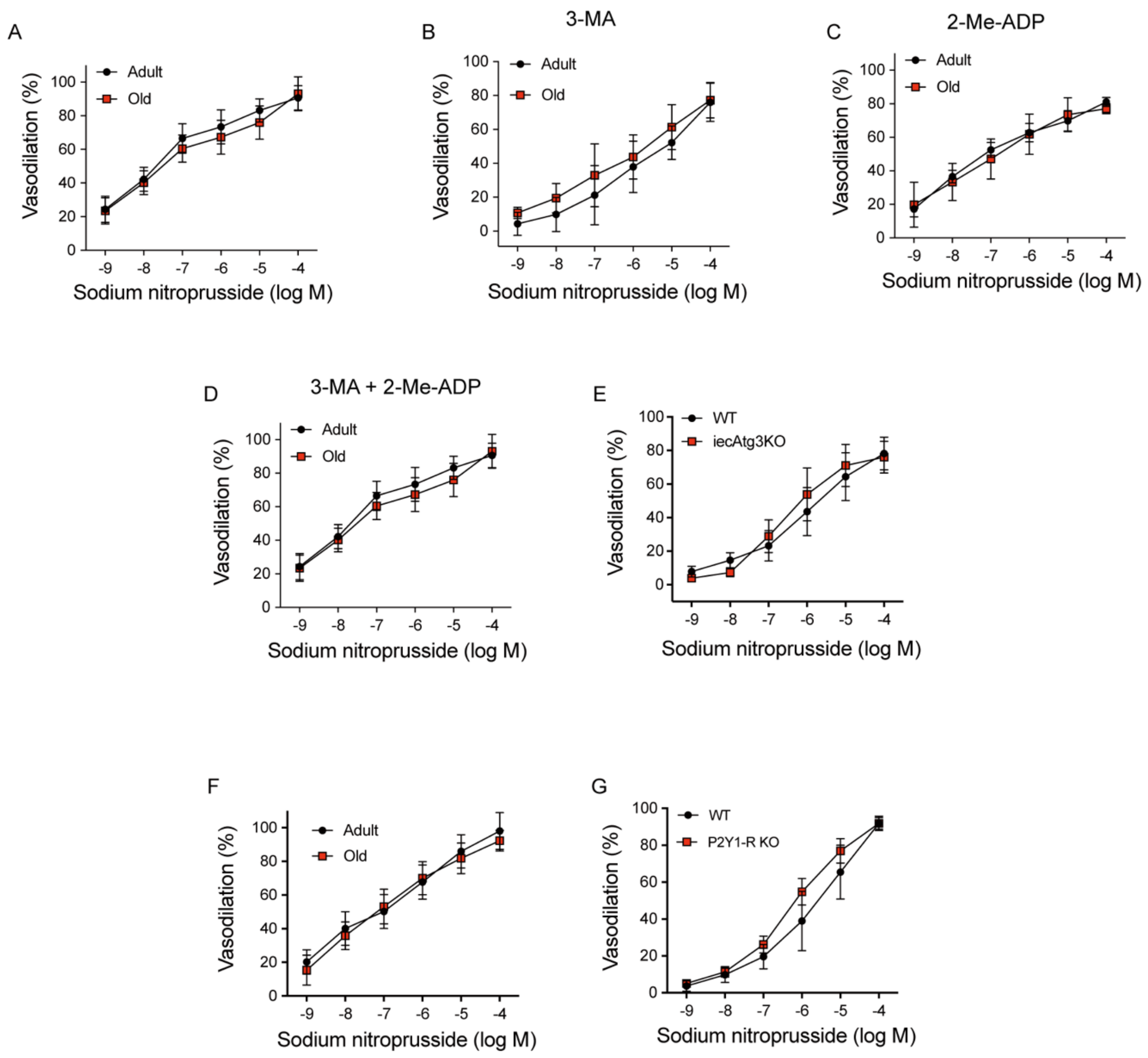

**Supplemental Figure VI. Responses to sodium nitroprusside in arteries.** Sodium nitroprusside evoked vasorelaxation in arteries isobarically pressurized to 60 mmHg was not different between adult and old mice (**A-D** and **F**), WT and iecAtg3KO mice (**E**), and WT vs $P2Y_1$-R KO mice (**G**). For **A**, n=4 mice per group. For **B**, n=4 mice per group. For **C-E**, n=5 mice per group. For **F**, n=3 mice per group. For **G**, n=4 mice per group. One artery per mouse was used. Statistical significance was assessed using a two-way repeated measures ANOVA.

HAECs

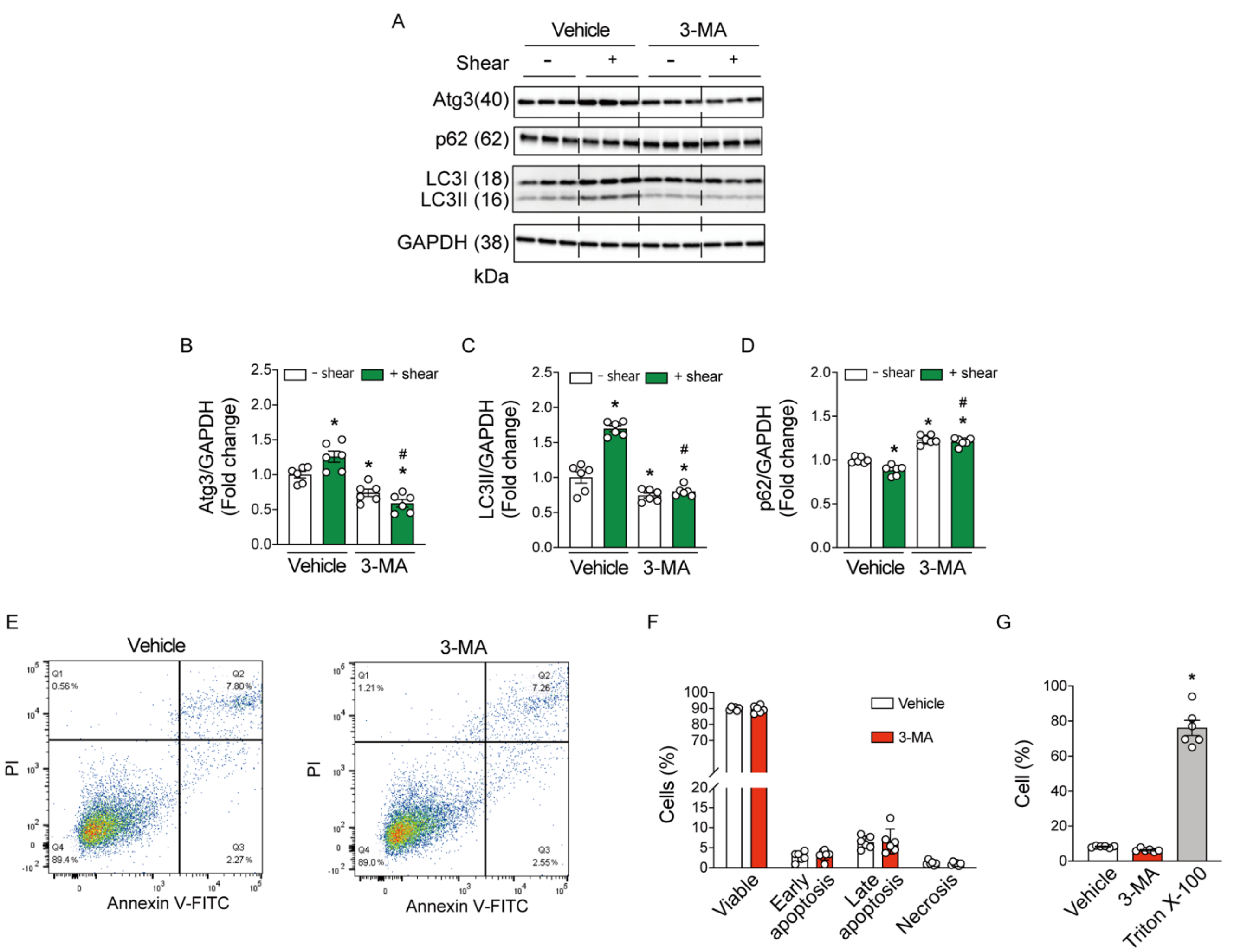

**Supplemental Figure VII. Shear stress-induced autophagy is prevented by 3-MA.** Human aortic endothelial cells (HAECs) treated with DMSO (vehicle) or 3-MA (5 mmol/L) were exposed to 0 dyne/cm2 (- shear) or 20 dyne/cm$^2$ (+ shear) for 45 min. Representative images (**A**) and mean densitometry (**B**) indicate Atg3:GAPDH and LC3II:GAPDH accumulation, and p62:GAPDH degradation, increased in HAECs exposed to 20 vs. 0 dyne/cm$^2$. Concurrent incubation with 3-MA prevented each response. For **B-D**, differences among groups concerning mean densitometry were identified using a one-way ANOVA; n=6 wells of a 6-well plate per treatment; *p<0.05 vs. vehicle. (**E**) Apoptosis was assessed among groups via Annexin V and propidium iodide (PI) staining. Representative images indicate necrosis (Quadrant 1; Q1); (ii) late apoptosis (Late, Q2); (iii) early apoptosis (Early, Q3); or (iv) viable cells (Q4). %, the percentage of cells in that particular quadrant. (**F**) Quantitative analysis of apoptosis indicates no difference between vehicle and 3-MA treated HAECs. (**G**) Cell death assessed using PrestoBlue indicated no differences between treatments, but a robust response to the positive control Triton-X-100. For **F** and **G**, differences were identified using a one-way ANOVA; n=6 wells of a 6-well plate per treatment; *p<0.05 vs. vehicle; #p<0.05 vs vehicle + Shear. Statistical significance was assessed using an unpaired t-test.

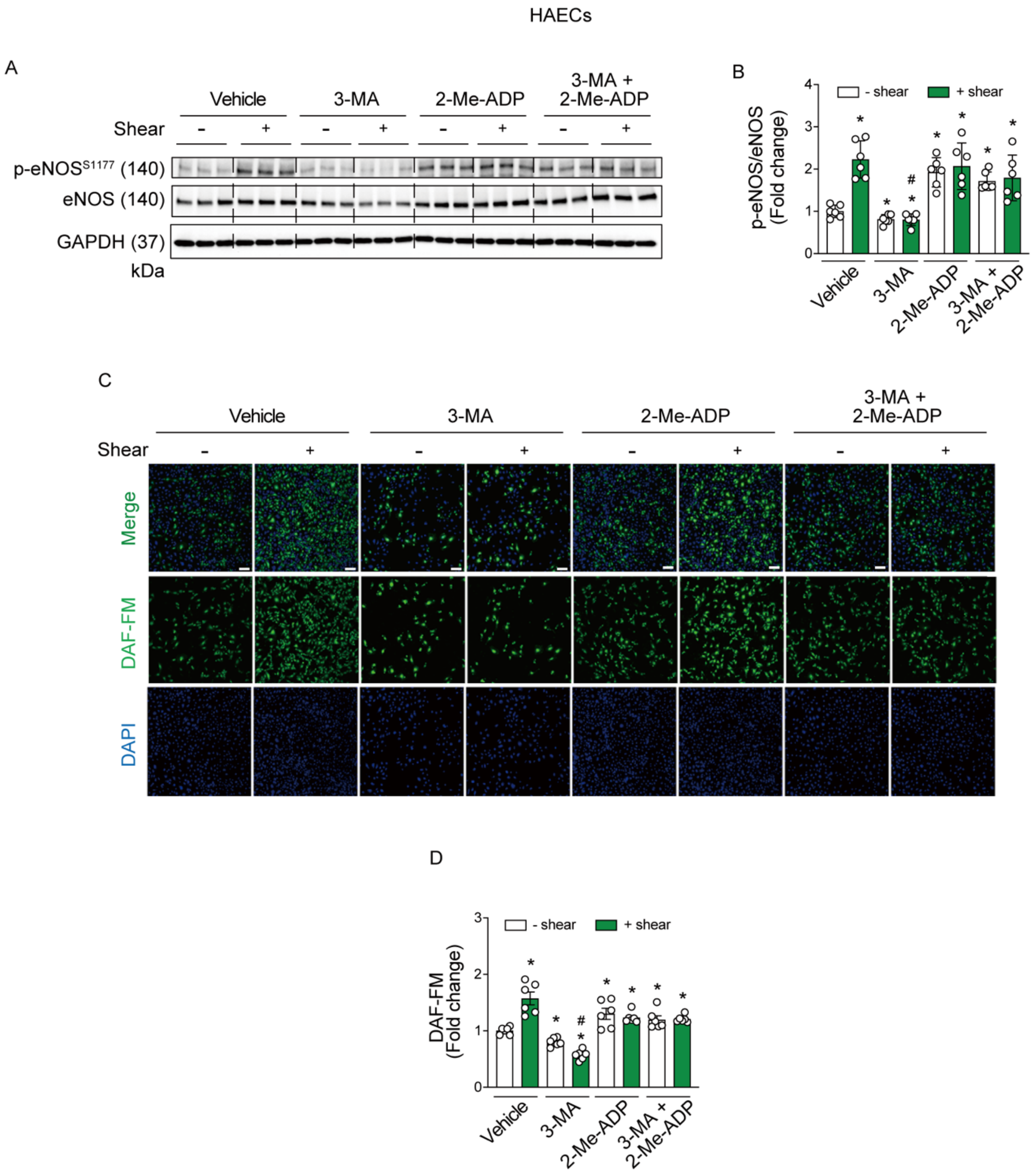

**Supplemental Figure VIII. Shear stress -induced eNOS activation and NO generation in HAECs is prevented by 3-MA but is restored by 2-Me-ADP.** HAECs treated with vehicle (DMSO), 3MA, 2-Me-ADP (100 µmol/L), or 3-MA + 2-Me-ADP were exposed to 0 (- shear) or 20 dyne/$cm^2$ shear stress (+shear). Representative images (**A** and **C**) and mean data (**B** and **D**) indicate shear-induced p-eNOS$^{S1177}$:eNOS and NO generation in the presence of vehicle was prevented by 3-MA, but is restored via concurrent incubation with 2-Me-ADP. For **B** (n=6) and **D** (n=5), differences among groups concerning mean densitometry or 4-amino-5methylamino-2',7'-difluorofluorescein diacetate (DAF-FM) fluorescence intensity, respectively, were identified using a one-way ANOVA; *p<0.05 vs. vehicle; #p<0.05 vs vehicle + Shear. Statistical significance was assessed using a one-way ANOVA.

HAECs

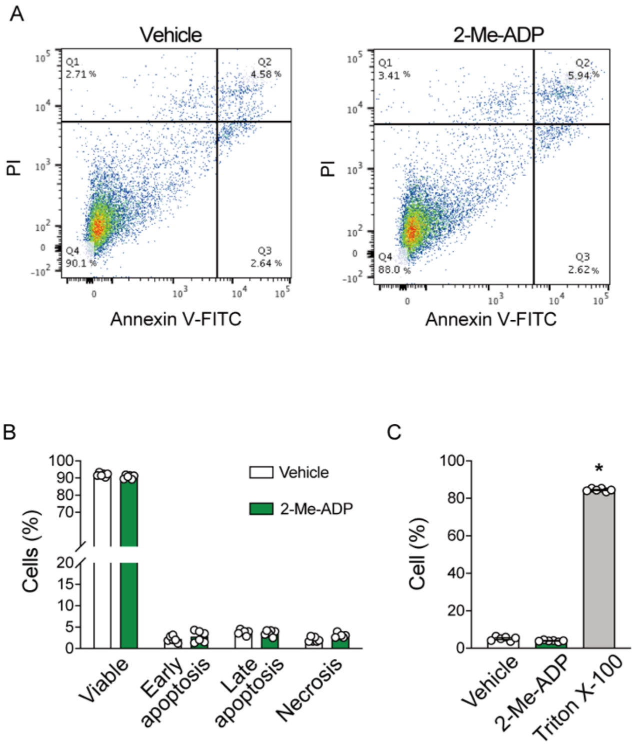

**Supplemental Figure IX. 2-Me-ADP treatment has no impact on apoptosis or cell death in HAECs.** (**A**) Apoptosis was assessed among groups via Annexin V and propidium iodide (PI) staining. Representative images indicate necrosis (Quadrant 1; Q1); (ii) late apoptosis (Late, Q2); (iii) early apoptosis (Early, Q3); or (iv) viable cells (Q4). %, the percentage of cells in that particular quadrant. (**B**) Quantitative analysis of apoptosis in vehicle and 2-Me-ADP treated HAECs. (**C**) Cell death assessed in vehicle and 2-Me-ADP treated HAECs using PrestoBlue. TritonX-100 was used as a positive control. For **B** and **C**, n=6 per treatment; *p<0.05 vs vehicle. Significance was assessed using an unpaired t-test (**B**) and a one-way ANOVA (**C**).

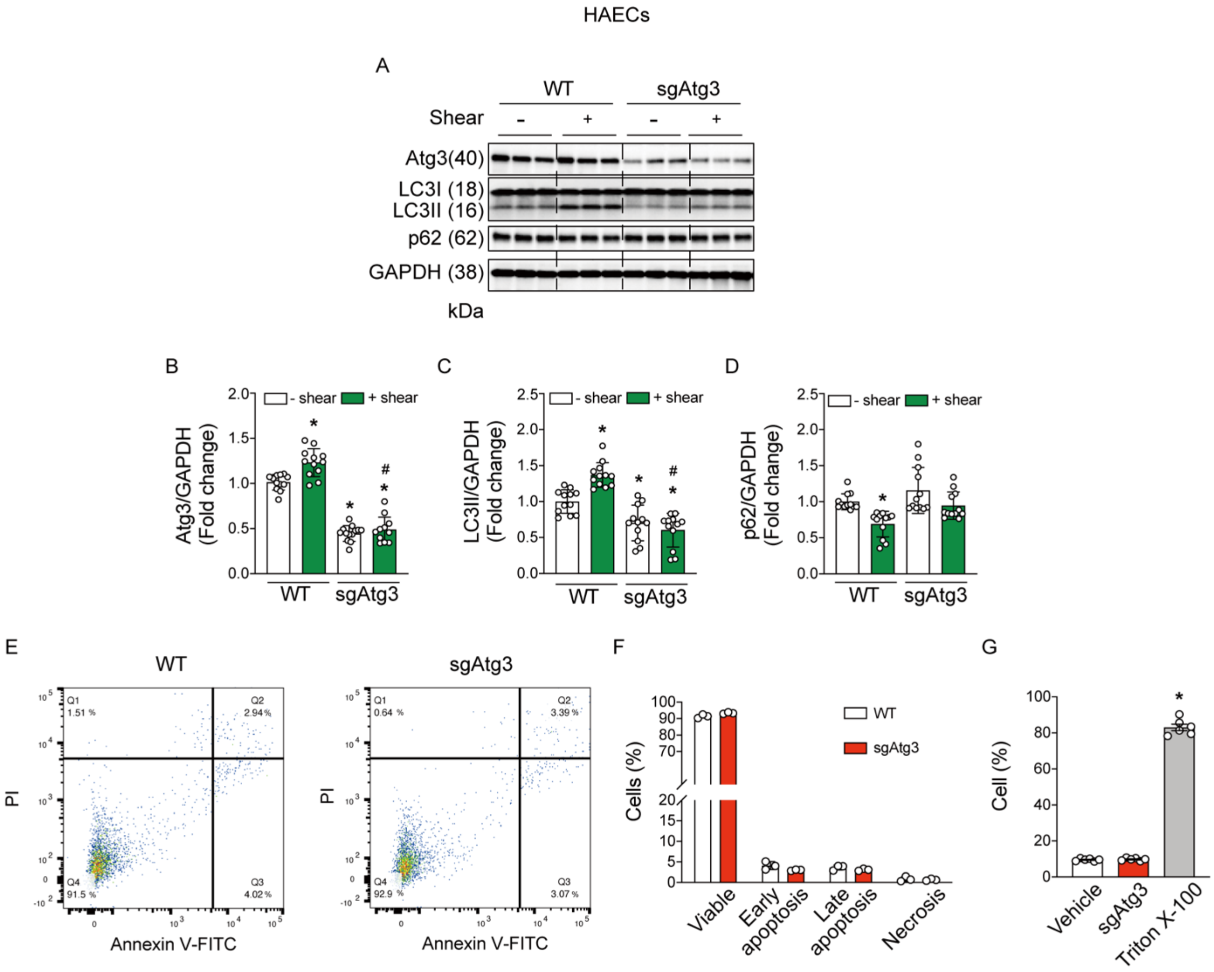

**Supplemental Figure X. Shear stress-induced autophagy is prevented in HAECs by depletion of Atg3 in the absence of cell death and apoptosis.** Atg3 was deleted in HAECs (sgAtg3) using CRISPR-Cas9. Next, HAECs were exposed to 0 (- shear) or 20 dyne/ $cm^2$ (+ shear) for 45-min. Representative images (**A**) and mean densitometry of Atg3 (**B**), LC3II (**C**), and p62 (**D**). For **B-D**, protein expression was normalized by GAPDH; n=12 for each condition. *p<0.05 vs WT (- shear); # vs. WT + shear. (**E**) Apoptosis was assessed among groups via Annexin V and propidium iodide (PI) staining. Representative images indicate necrosis (Quadrant 1; Q1); (ii) late apoptosis (Late, Q2); (iii) early apoptosis (Early, Q3); or (iv) viable cells (Q4). %, the percentage of cells in that particular quadrant. (**F**) Quantitative analysis of apoptosis in WT and sgAtg3 HAECs. (**G**) Cell death assessed in WT and sgAtg3 HAECs using PrestoBlue. TritonX-100 was used as a positive control. For **B** and **C**, n=6 per treatment; *p<0.05 vs WT. Statistical significance was assessed using a one-way ANOVA (B-D, G) and by an unpaired t-test (**F**).

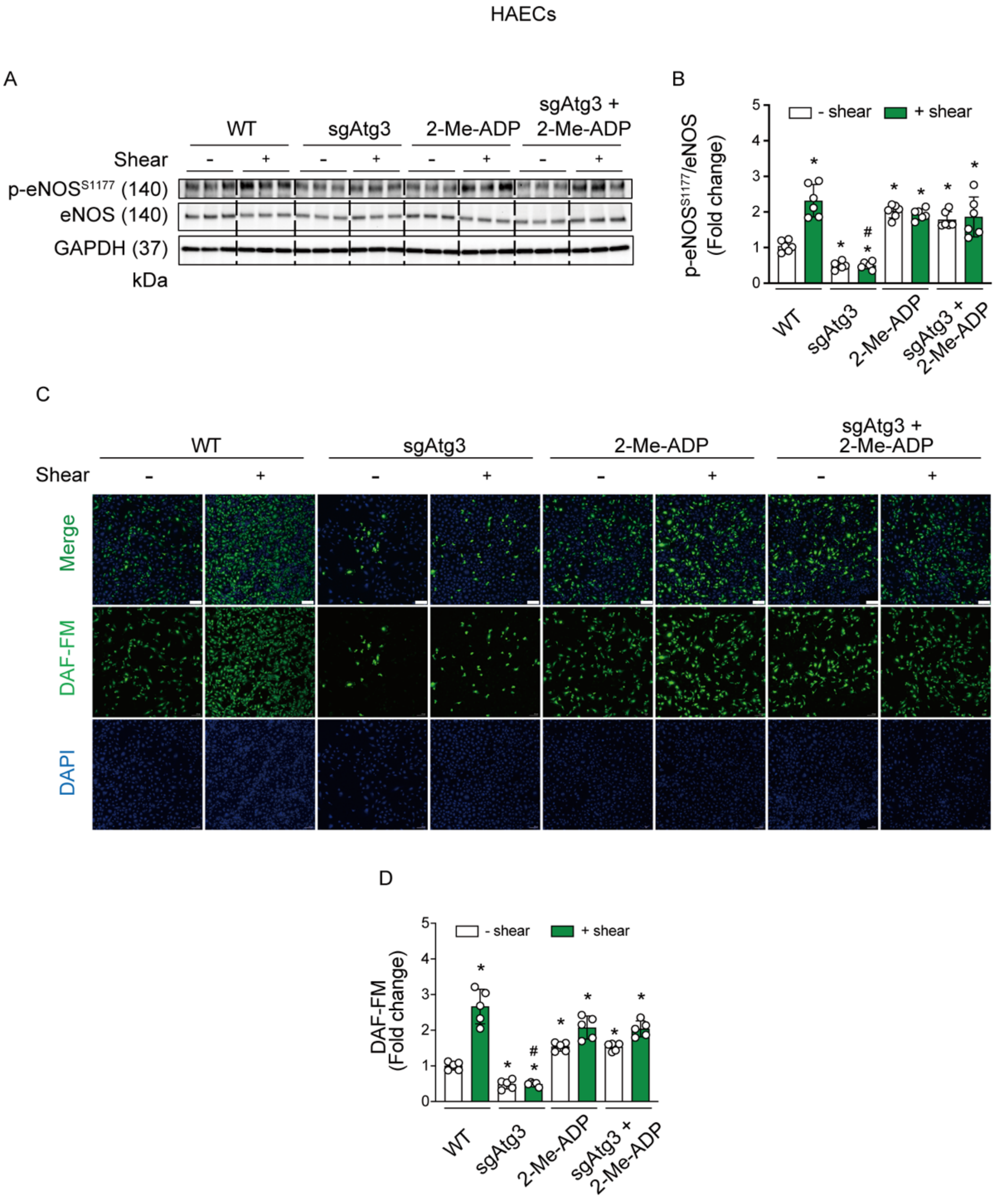

**Supplemental Figure XI. Shear stress-induced eNOS activation and NO generation in HAECs is prevented by Atg3 knockdown but is restored by 2-Me-ADP**. Representative images (**A** and **C**) and mean densitometry of p-eNOS$^{S1177}$ : eNOS protein expression (**B**) and DAF-FM staining intensity (**D**; n=6 wells of a 6-well plate per condition for both). Compared to 0 dyne / cm$^2$ (- shear), 20 dyn / cm$^2$ for 45-min (+ shear) increased p-eNOS$^{S1177}$ protein expression and DAF-FM staining intensity in WT HAECs (histogram 1 vs. 2) but not HAECs transfected with sgAtg3 (histogram 3 vs. 4). Treatment with 100 µmol/L 2-Me-ADP restored shear-induced p-eNOS$^{S1177}$ and DAF-FM staining intensity in sgAtg3 HAECS to levels not different from WT HAECs (histogram 2 vs. 8). For **C**, scale bar represents 20 µm, magnification=10X. *$p<0.05$ vs WT-shear; # vs. WT+shear. Statistical significance was assessed using a two-way ANOVA.

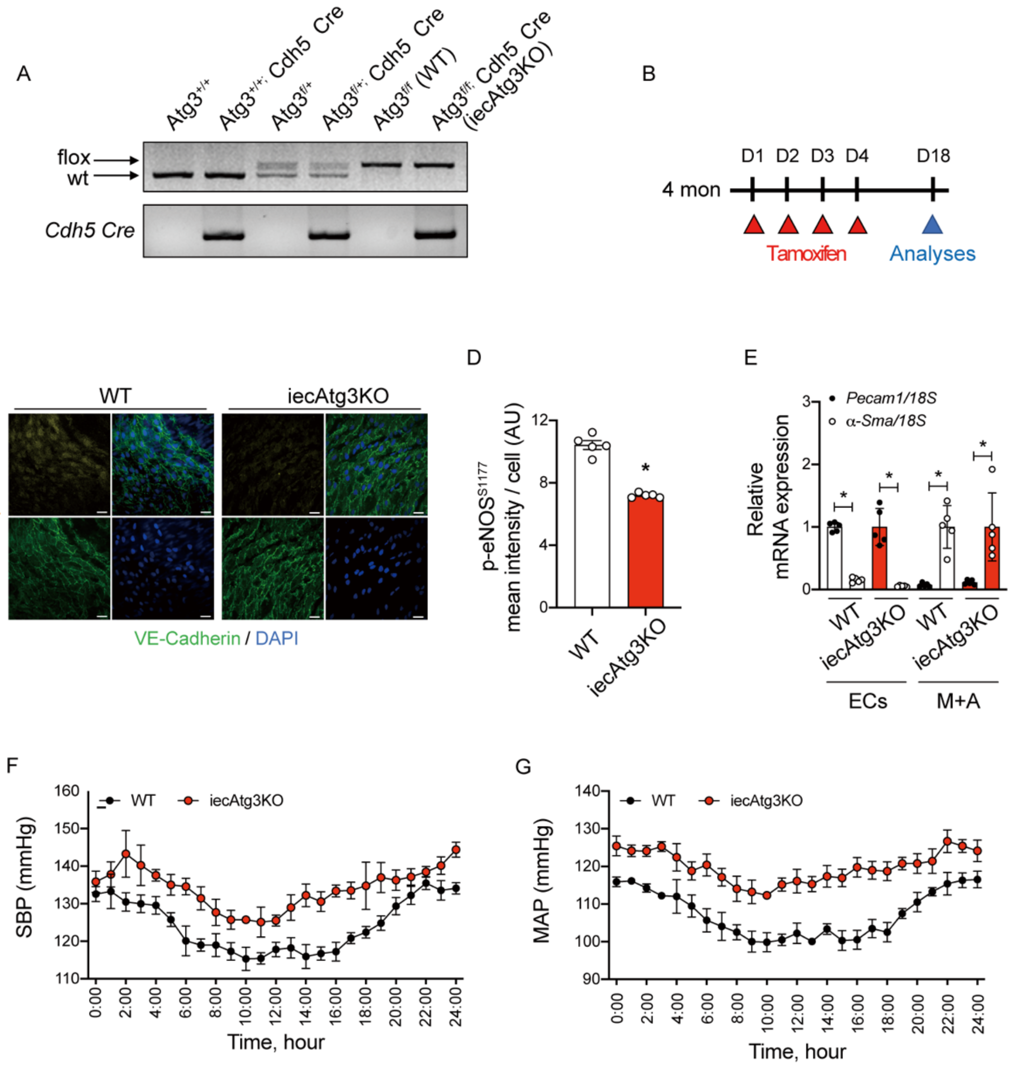

**Supplemental Figure XII. Mice with tamoxifen-inducible endothelial cell specific Atg3 depletion.** (**A**) Representative image of genotyping of Atg3 flox and Cdh5 (VE-Cadherin) Cre mice. Atg3 flox/flox mice were used as wild type (WT) mice and Atg3 flox/flox and Cdh5 Cre positive mice were used as iecAtg3KO mice. (**B**) To activate the Cre-LoxP system, tamoxifen (4 mg) was administered for 4 consecutive days to 4 month old WT and iecAtg3KO mice. Experiments were initiated 14 d after the last tamoxifen injection. Representative *en face* images (**C**) and mean immunofluorescent staining intensity (normalized by the number of cells; (**D**) of endothelial cells (ECs) from aorta of wild type (WT) mice and mice with inducible depletion of Atg3 in ECs. ECs were identified by co-staining for DAPI and VE-cadherin. For each quadrant (**C**): bottom right DAPI (blue); bottom left, VE-cadherin (red); top left: p-eNOS$^{S1177}$ (yellow); top right, merge. p-eNOS$^{S1177}$ (**D**) was lower in ECs from iecAtg3KO vs. WT mice. For **C** and **D**, n=3 segments of aorta from 4 mice in each group. For *C*, scale bar represents 10 μm, magnification=60X. (**E**) qPCR was used to assess mRNA expression in the endothelial cell (EC) and media + adventitia (M+A) fraction obtained from carotid arteries of WT and iecAtg3KO mice. *Pecam1* mRNA expression is robust whereas *α-Sma* is minimal in the EC fraction obtained from carotid arteries of WT and iecAtg3KO mice. Conversely, *α-Sma* mRNA expression is robust whereas *Pecam1* is minimal in the M+A fraction obtained from carotid arteries of WT and iecAtg3KO mice. Data shown in *E* were normalized by *18S*; n=5 mice per group x 2 carotid arteries per mouse. For **E**, *p<0.05 vs *Pecam1*. SBP (**F**) and MAP (**G**) data represent hourly averages obtained during 3 x 24 h periods obtained 6-days post surgery. These data indicate blood pressure is greater in iecAtg3KO vs. WT mice. For **F** and **G**, n=6 per group. *p<0.05 vs WT. Statistical significance was assessed by unpaired t-test (**A** and **B**).

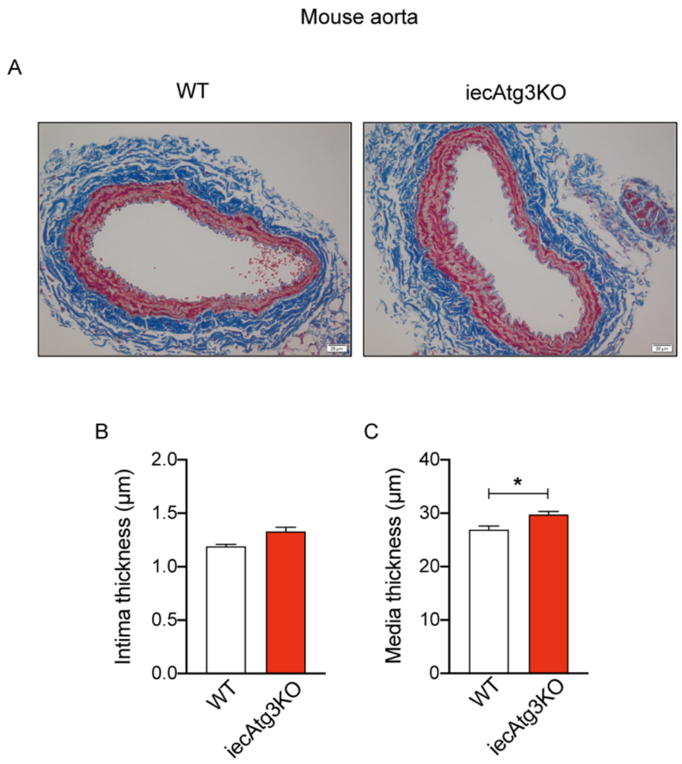

**Figure XIII. Media thickness is greater in aorta from iecAtg3KO vs. WT mice.** A representative image of trichrome staining (**A**) and mean data concerning: (**B**) intima thickness; and (**C**) media thickness. Scale bar represents 20 µmol/L. Magnification=10X. For **B** and **C**, n=5 mice, one segment of aorta per mouse, obtained just proximal to the bifurcation of the iliac arteries. *p<0.05 vs WT. Statistical significance was assessed using an unpaired t-test.

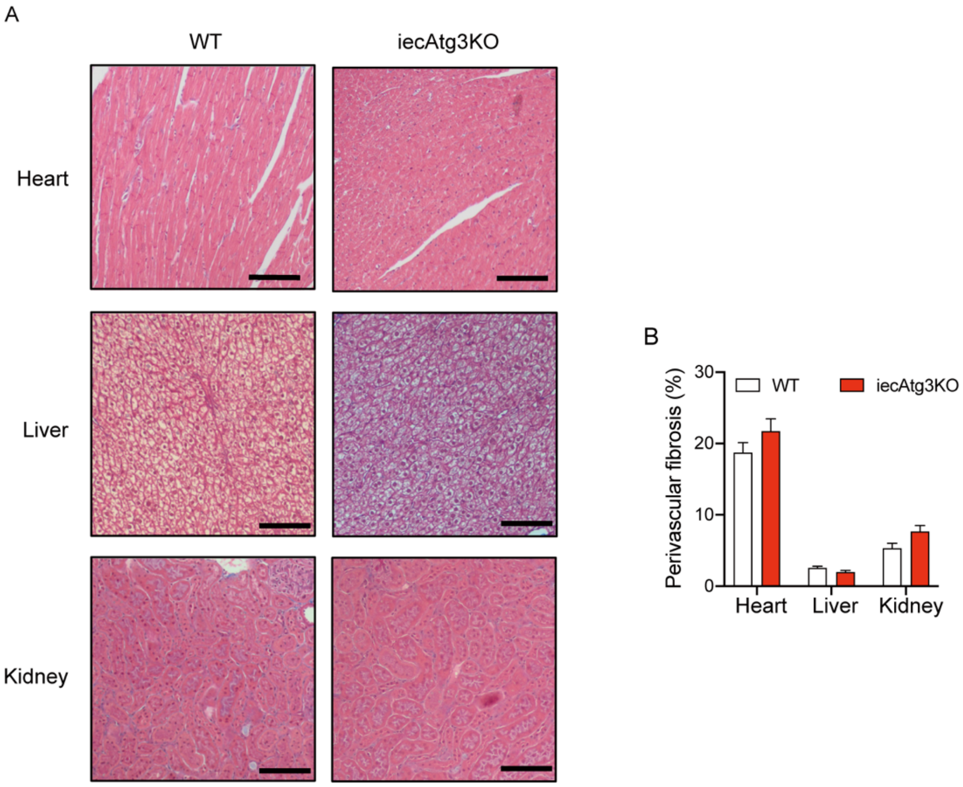

**Figure XIV. Perivascular fibrosis displayed by heart, liver, and kidney is not different between WT and iecAtg3KO mice.** (**A**) Representative images of trichrome staining in heart, liver, and kidney from WT and iecAtg3KO mice. (**B**) Collagen was assessed to determine perivascular fibrosis in heart, liver, and kidney from WT and iecAtg3KO mice. For A, the scale bar represents 20 μm. For **B**, n=3 mice from each group. Magnification=10X. Statistical significance was assessed using an unpaired t-test.

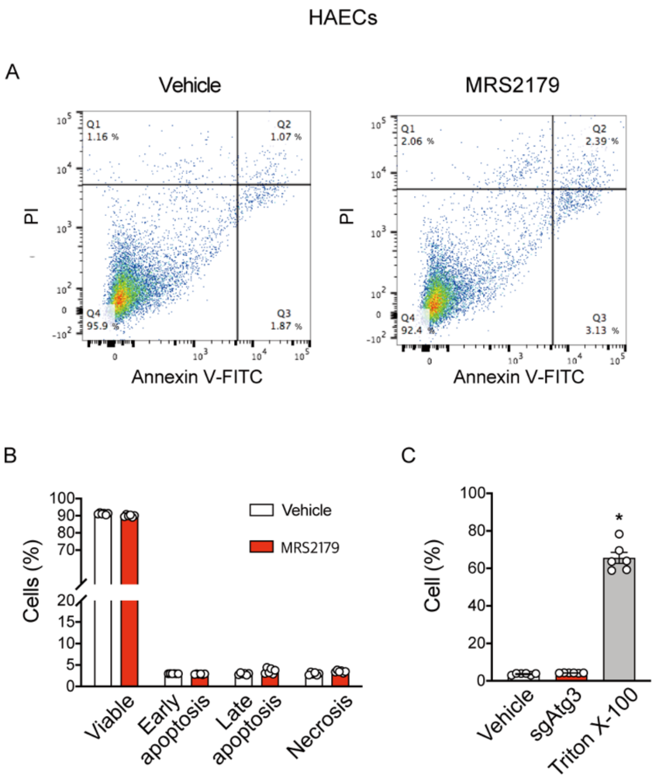

**Figure XV. P2Y$_1$-R blockade using MRS2179 has no impact on apoptosis or cell death in HAECs**. (**A**) Apoptosis was assessed among groups via Annexin V and propidium iodide (PI) staining. Representative images indicate necrosis (Quadrant 1; Q1); (ii) late apoptosis (Late, Q2); (iii) early apoptosis (Early, Q3); or (iv) viable cells (Q4). %, the percentage of cells in that particular quadrant. (**B**) Quantitative analysis of apoptosis in vehicle and MRS2179 treated HAECs. (**C**) Cell death assessed in vehicle and MRS2179 treated HAECs using PrestoBlue. TritonX-100 was used as a positive control. For **B** and **C**, n=6 per treatment; *p<0.05 vs vehicle. Statistical significance was assessed using an unpaired t-test (**B**) and a one-way ANOVA (**C**).

**Supplemental Tables**

**Supplemental Table I. Characteristics of adult and old human subjects**

| | Adult | Old |
|---|---|---|
| Age, years | 23 ± 1 | 68 ± 2* |
| Height, cm | 181 ± 3 | 180 ± 2 |
| Weight, kg | 75 ± 2 | 91 ± 9 |
| Body mass index, $kg/m^2$ | 23 ± 1 | 28 ± 2* |
| Cholesterol, mg/dl | 137 ± 16 | 149 ± 22 |
| Triglycerides, mg/dl | 50 ± 5 | 111 ± 31 |
| HDL, mg/dl | 45 ± 3 | 41 ± 6 |
| LDL, mg/dl | 83 ± 13 | 94 ± 21 |
| Glucose, mg/dl | 81 ± 2 | 83 ± 7 |
| RBC, M/uL | 5 ± 0 | 5 ± 0 |
| Hb, g/dl | 15 ± 0 | 15 ± 0 |
| Hct % | 42 ± 1 | 44 ± 1 |
| Maximal workload, kg | 25 ± 2 | 19 ± 1 |

HDL, high-density lipoproteins; LDL, low-density lipoproteins; RBC, red blood cells; Hb, hemoglobin; Hct, hematocrit. n=6 per group. Values are means ± SE. *p<0.05 vs. Adult using an unpaired t-test.

**Table S2. Hemodynamics during rhythmic handgrip exercise**

| Adult | Baseline | 5 min | 10 min | 15 min | 20 min | 30 min | 40 min | 50 min | 60 min |
|---|---|---|---|---|---|---|---|---|---|
| Brachial artery velocity, cm/s | 7.9 ± 0.8 | 17.1 ± 1* | 19.8 ± 2.1* | 20.6 ± 1.8* | 20.8 ± 1.6* | 21.1 ± 1.1* | 21.2 ± 1.4* | 18.9 ± 0.8* | 20.2 ± 1.2* |
| Heart rate, bpm | 56 ± 1 | 60 ± 1 | 61 ± 1 | 62 ± 2 | 60 ± 2 | 60 ± 1 | 60 ± 2 | 61 ± 1 | 59 ± 1 |
| Stroke volume, ml | 102 ± 5 | 101 ± 5 | 103 ± 5 | 103 ± 5 | 105 ± 5 | 105 ± 5 | 104 ± 5 | 107 ± 6 | 105 ± 6 |
| Cardiac output, L/min | 5.7 ± 0.3 | 6.2 ± 0.4 | 6.3 ± 0.3 | 6.3 ± 0.3 | 6.3 ± 0.4 | 6.3 ± 0.4 | 6.2 ± 0.4 | 6.6 ± 0.5 | 6.2 ± 0.4 |
| Mean arterial pressure, mmHg | 90 ± 3 | 9 ± 2 | 93 ± 2 | 92 ± 2 | 95 ± 2 | 94 ± 2 | 93 ± 2 | 96 ± 2 | 97 ± 2 |
| Rate of perceived exertion, rating | | 1.8 ± 0.6 | 3.1 ± 0.4 | 3.1 ± 0.4 | 3.2 ± 0.4 | 3.3 ± 0.4 | 3.3 ± 0.4 | 3.2 ± 0.3 | 3.3 ± 0.3 |
| **Old** | | | | | | | | | |
| | Baseline | 5 min | 10 min | 15 min | 20 min | 30 min | 40 min | 50 min | 60 min |
| Brachial artery velocity, cm/s | 7.6 ± 0.9 | 17.5 ± 2.5* | 17.3 ± 2.4* | 18.5 ± 2.3* | 18.3 ± 2.4* | 18.9 ± 1.9* | 19.2 ± 1.8* | 20.2 ± 2* | 20.0 ± 2.2* |
| Heart rate, bpm | 49 ± 2# | 53 ± 4 | 51 ± 3# | 53 ± 3 | 53 ± 3 | 54 ± 3 | 53 ± 3 | 54 ± 3 | 52 ± 3 |
| Stroke volume, ml | 102 ± 13 | 99 ± 12 | 100 ± 10 | 102 ± 10 | 101 ± 10 | 98 ± 10 | 96 ± 14 | 87 ± 12 | 92 ± 13 |
| Cardiac output, L/min | 5.1 ± 0.8 | 5.5 ± 0.9 | 5.5 ± 0.7 | 5.6 # 0.8 | 5.6 ± 0.8 | 5.4 ± 0.7 | 5.3 ± 0.9 | 4.9 ± 0.8 | 5.1 ± 0.8 |
| Mean arterial pressure, mmHg | 102 ± 6 | 106 ± 6 | 111 ± 6# | 109 ± 7# | 111 ± 6# | 113 ± 6# | 112 ± 6# | 118 ± 5# | 115 ± 5# |
| Rate of perceived exertion, rating | | 2.9 ± 0.5 | 4.0 ± 0.7 | 4.6 ± 0.9 | 5.3 ± 1.0 | 6.0 ± 1# | 6.6 ± 0.9# | 6.3 ± 0.8# | 5.9 ± 1# |

BA, brachial artery; HR, heart rate; SV, stroke volume; CO, cardiac output; MAP, mean arterial pressure; RPE, rate of perceived exertion. Note : 0-min = RHE-Pre; 60-min = RHE-Post. n=6 per group. Values are means ± SE. *$p<0.05$ vs. 0-min; # $p<0.0.5$ vs adult using a two-way repeated measures ANOVA.

**Supplemental Table III. Characteristics of adult and old mice**

| | Adult | Old |
|---|---|---|
| *Animal characteristics* | | |
| Age, months | 7 ± 1 | 23 ± 1* |
| Blood glucose, mg/dl | 135 ± 18 | 131 ± 16 |
| Body weight, g | 30 ± 1 | 32 ± 1 |
| *Femoral a. characteristics* | | |
| Diameter at 0 mmHg, μm | 216 ± 7 | 229 ± 12 |
| Diameter at 60 mmHg, μm | 319 ± 13 | 324 ± 14 |

Blood glucose was obtained via tail snip from random-fed mice. Diameter, external diameter. n=6 per group. Values are means ± SE.

**Supplemental Table IV. Characteristics of adult and old mice treated with 3-MA, 2-Me-ADP, or 3-MA+2-Me-ADP**

| ***3-MA*** | Adult | Old |
|---|---|---|
| *Animal characteristics* | | |
| Age, months | 7 ± 1 | 23 ± 1* |
| Body weight, g | 33 ± 1 | 36 ± 1 |
| *Femoral a. characteristics* | | |
| Diameter at 0 mmHg, µm | 215 ± 4 | 218 ± 6 |
| Diameter at 60 mmHg, µm | 298 ± 5 | 304 ± 5 |
| ***2-Me-ADP*** | Adult | Old |
| *Animal characteristics* | | |
| Age, months | 7 ± 1 | 23 ± 1 |
| Body weight, g | 33 ± 1 | 37 ± 1 |
| *Femoral a. characteristics* | | |
| Diameter at 0 mmHg, µm | 191 ± 4 | 196 ± 3 |
| Diameter at 60 mmHg, µm | 304 ± 4 | 300 ± 3 |
| ***3-MA + 2-Me-ADP*** | Adult | Old |
| *Animal characteristics* | | |
| Age, months | 7 ± 1 | 23 ± 1 |
| Body weight, g | 34 ± 1 | 38 ± 3 |
| *Femoral a. characteristics* | | |
| Diameter at 0 mmHg, µm | 226 ± 12 | 246 ± 12 |
| Diameter at 60 mmHg, µm | 346 ± 16 | 353 ± 10 |

For 3-MA, n=4 for each group; For 2-Me-ADP, n=5 of each group; For 3-MA + 2-Me-ADP, n=4-5. Values are means ± SE. Statistical significance was assessed by unpaired t-test.

**Supplemental Table V. Characteristics of wild type (WT) animals and mice with inducible depletion of Atg3 specifically in endothelial cells (iecAtg3KO mice)**

| | WT | iecAtg3KO |
|---|---|---|
| *Animal characteristics* | | |
| Age, months | 4 ± 1 | 4 ± 1 |
| Blood glucose, mg/dl | 158 ± 19 | 151 ± 24 |
| Body weight, g | 26 ± 1 | 28 ± 1 |
| *Femoral a. characteristics* | | |
| Diameter at 0 mmHg, µm | 257 ± 25 | 225 ± 9 |
| Diameter at 60 mmHg, µm | 361 ± 19 | 331 ± 19 |

Diameter, external diameter. n=5 per group. Values are means ± SE.

**Supplemental Table VI. Characteristics of adult and old mice treated with MRS2179 and of $P2Y_1$-R KO mice**

| ***MRS2179*** | Adult | Old |
|---|---|---|
| *Animal characteristics* | | |
| Age, months | 7 ± 1 | 23 ± 1* |
| Body weight, g | 32 ± 1 | 37 ± 2 |
| *Femoral a. characteristics* | | |
| Diameter at 0 mmHg, µm | 217 ± 3 | 215 ± 3 |
| Diameter at 60 mmHg, µm | 304 ± 15 | 286 ± 3 |
| | WT | $P2Y_1$-R KO |
| *Animal characteristics* | | |
| Age, months | 2 ± 1 | 2 ± 1 |
| Body weight, g | 22 ± 1 | 23 ± 1 |
| *Femoral a. characteristics* | | |
| Diameter at 0 mmHg, µm | 241 ± 8 | 217 ± 11 |
| Diameter at 60 mmHg, µm | 341 ± 15 | 327 ± 12 |

Diameter, external diameter. For MRS2179, n=3 per group; for $P2Y_1$-R KO mice, N=4 per group. Values are means ± SE. Statistical significance was assessed by unpaired t-test.

**Supplemental Table VII.** Primers used for genotyping

| Name | Forward Primer sequence (5'-3') | Reverse Primer sequence (5'-3') |
|---|---|---|
| *Atg3 flox* | GTACCTGACCCCGGTCCT | TTGGACAGTGGACTAAGTG |
| *Cdh5 Cre* | CTGCTGGGATGCTGAAGGCATC | TGTCCATCAGGTTCTTGCGAACC |

**Supplemental Table VIII. Primers used for RT-qPCR**

| Gene name | Forward Primer sequence (5'-3') | Reverse Primer sequence (5'-3') | NCBI Gene ID | Application size (bp) |
|---|---|---|---|---|
| *Pecam1* | CCAAGGCCAAACAGA | AAGGGAGCCTTCCGTTCT | 18613 | 323 |
| *α-Sma* | CAGGGAGTAATGGTTGGAAT | TCTCAAACATAATCTGGGTCA | 11475 | 256 |
| *Tfeb* | CCACCCCAGCCATCAACAC | CAGACAGATACTCCCGAACCTT | 21425 | 147 |
| *Beclin1* | ATGGAGGGGTCTAAGGCGTC | TCCTCTCCTGAGTTAGCCTCT | 56208 | 197 |
| *Atg3* | ACACGGTGAAGGGAAAGGC | TGGTGGACTAAGTGATCTCCAG | 67841 | 130 |
| *Atg5* | TGTGCTTCGAGATGTGTGGTT | GTCAAATAGCTGACTCTTGGCAA | 11793 | 120 |
| *Atg7* | GTTCGCCCCCTTTAATAGTGC | TGAACTCCAACGTCAAGCGG | 11793 | 161 |
| *Map1lc3b* | TTATAGAGCGATACAAGGGGGAG | CGCCGTCTGATTATCTTGATGAG | 67443 | 109 |
| *Sqstm1* | ATGTGGAACATGGAGGGAAGA | GGAGTTCACCTGTAGATGGGT | 18412 | 178 |
| *B2m* | TTCTGGTGCTTGTCTCACTGA | CAGTATGTTCGGCTTCCCATTC | 12010 | 104 |
| *Hprt* | TCAGTCAACGGGGGACATAAA | GGGGCTGTACTGCTTAACCAG | 15452 | 142 |
| *Actb* | GGCTGTATTCCCCTCCATCG | CCAGTTGGTAACAATGCCATGT | 11461 | 154 |
| *18S* | GTAACCCGTTGAACCCCATT | CCATCCAATCGGTAGTAGCG | 19791 | 151 |

### Major Resources Table

**Animals (in vivo studies)**

| Species | Vendor or Source | Background Strain | Sex | Persistent ID / URL |
|---|---|---|---|---|
| Mouse | JAX lab | C57BL/6J | Male | |
| Mouse | NIA | C57BL/6J | Male | |

**Genetically Modified Animals**

| | Species | Vendor or Source | Background Strain | Other Information | Persistent ID / URL |
|---|---|---|---|---|---|
| Atg3 flox/flox | Mouse | University of Utah Transgenic and Mouse Core Facility | C57BL/6J | | |
| Cdh5-Cre$^{ERT2}$ | Mouse | University of Munster | C57BL/6J | Dr. Ralf Adams provided | |
| $P2Y_1$-R KO | Mouse | Temple University | C57BL/6J | Dr. Satya Kunapuli provided | |

**Antibodies**

| Target antigen | Vendor or Source | Catalog # | Working concentration | Lot # (preferred but not required) | Persistent ID / URL |
|---|---|---|---|---|---|
| p-eNOS$^{S1177}$ (Human, mouse) | Thermo Fisher Scientific | PA5-35879 | 1:100 (IF)<br>1:1,000 (IB) | | |
| eNOS (Human) | BD Bioscience | 610297 | 1:100 (IF) | | |
| eNOS (Human, Mouse) | Cell Signaling | 32027 | 1:1,000 (IB) | | |
| Beclin-1 (Human) | Novus Biologicals | NB500-249 | 1:100 (IF) | | |
| Atg3 (Human, Mouse) | Thermo Fisher Scientific | MA5-25610 | 1:100 (IF) | | |
| Atg3 (Human) | Abcam | ab108251 | 1:1,000 (IB) | | |
| LC3B (Human, Mouse) | Cell signaling | 83506S | 1:200 (IF) | | |
| LC3B (Human) | Sigma Aldrich | L7543 | 1:500 (IB) | | |
| P62 (Human) | Abcam | ab56416 | 1:100 (IF)<br>1:1,200 (IB) | | |

| | | | |
|---|---|---|---|
| P62 (Mouse) | Abnova | H00008878-M01 | 1:100 (IF) |
| LAMP2A (Human) | Abcam | ab18528 | 1:100 (IF) |
| VE-Cadherin (Human) | R&D System | AF938 | 1:30 (IF) |
| VE-Cadherin (Mouse) | R&D System | AF1002 | 1:30 (IF) |
| GAPDH (Human, Mouse) | Cell Signaling | 2118 | 1:1,200 (IB) |
| Donkey anti-goat IgG H&L (Alexa Fluor 488) | Abcam | ab150129 | 1:500 (IF) |
| Donkey anti-mouse IgG H&L (Alexa Fluor 488) | Abcam | ab150109 | 1:500 (IF) |
| Donkey anti-rabbit IgG H&L (Alexa Fluor 568) | Abcam | ab175692 | 1:400 (IF) |
| Donkey anti-mouse IgG H&L (Alexa Fluor 647) | Abcam | ab150107 | 1:250 (IF) |
| Donkey anti-goat IgG H&L (Alexa Fluor 647) | Abcam | ab150135 | 1:250 (IF) |
| Goat anti-Rabbit | Thermo Fisher Scientific | 31460 | 1:5,000 |
| Goat anti-Mouse | Thermo Fisher Scientific | 31430 | 1:10,000 |

**DNA/cDNA Clones (None)**

| Clone Name | Sequence | Source / Repository | Persistent ID / URL |
|---|---|---|---|

**Cultured Cells**

| Name | Vendor or Source | Sex (F, M, or unknown) | Persistent ID / URL |
|---|---|---|---|
| HAECs | Lonza Inc | | CC-2535 |

**Data & Code Availability (None)**

| Description | Source / Repository | Persistent ID / URL |
|---|---|---|

**Other**

| Description | Source / Repository | Persistent ID / URL |
|---|---|---|
| BSA | Sigma Aldrich | A1470 |
| 0.5 M EDTA, pH 8.0 | Thermo Fisher Scientific | 15575020 |
| Heparin | Sigma Aldrich | H3149 |
| Erythrocyte lysing kit | R&D system | WL1000 |
| EBM-2 Basal Medium | Lonza Inc | CC-3156 |
| EGM-2 SingleQuots Supplements | Lonza Inc | CC-4176 |
| Poly-L-lysine solution | Sigma Aldrich | P4707 |
| 4-Amino-5-Methylamino-2',7'-Difluorofluorescein Diacetate (DAF-FM Diacetate) | Thermo Fisher Scientific | D23844 |
| Dihydroethidium (DHE) | Thermo Fisher Scientific | D11347 |
| 4',6-Diamidino-2-Phenylindole Dihydrochloride (DAPI) | Thermo Fisher Scientific | D1306 |
| N-acetyl-L-cysteine | Sigma Aldrich | A7250 |
| $N^G$-Methyl-L-arginine acetate salt (L-NMMA) | Sigma Aldrich | M7033 |
| Paraformaldehyde | Sigma Aldrich | 158127 |
| Triton X-100 | Sigma Aldrich | X100 |
| ProLong Diamond Antifade mount with DAPI | Thermo Fisher Scientific | P36962 |
| 3-methyladenine (3-MA) | Sigma Aldrich | M9281 |
| 2-methylthioadenosine diphosphate trisodium salt (2-Me-ADP) | Tocris | 1624 |
| MRS2179 tetrasodium salt | Tocris | 0900 |
| Adenosine 5'-diphosphate sodium salt (ADP) | Sigma Aldrich | A2754 |
| PrestoBlue cell viability reagent | Thermo Fisher Scientific | A13261 |
| FITC Annexin V/Dead Cell Apoptosis Kit | Thermo Fisher Scientific | V13242 |
| SYBR green fluorescence | Qiagen | 204143 |
| Normal goat serum | Abcam | ab7481 |
| RIPA Lysis and Extraction buffer | Thermo Fisher Scientific | 89900 |

| | | |
|---|---|---|
| Laemmli SDS sample buffer | Alfa Aesar | J61337 |
| Blotting-Grade Blocker | Bio Rad | 1706404 |
| SuperSignal West Dura Extended Duration Substrate | Thermo Fisher Scientific | 34076 |

#### Supplemental Video Legend

**Supplemental Video I.** *Rhythmic handgrip exercise to elevate radial artery shear rate.* This procedure was completed during visit 2. Subjects were instrumented as described in the text. Radial artery endothelial cells (ECs) were obtained via j-wire before rhythmic handgrip exercise (i.e., RHE-Pre). Next, subjects squeezed and released a handle that elevated a bucket containing 5-10% of the weight achieved during their maximal handgrip workload test completed during visit 1. The duty cycle was 1 : 2 and subjects contracted to the sound of a metronome. The weight was designed to provide a workload that required ~ 3-fold elevations in radial artery shear rate for 60-min. The elevation of radial artery shear rate was estimated throughout the 60-min protocol by directly measuring brachial artery (BA) shear rate (see online methods) over 6 cardiac cycles at 10 min intervals. Rating of perceived exertion (RPE) was assessed according to the modified 10 point Borg scale i.e., mild (0-3), moderate (4-7), or difficult (8-10). Small weights were added or taken out of the bucket in an effort to maintain 3-fold elevations in arterial shear- rate.
